# Signatures of rapid synaptic learning in the hippocampus during novel experiences

**DOI:** 10.1101/2021.07.02.450956

**Authors:** James B. Priestley, John C. Bowler, Sebi V. Rolotti, Stefano Fusi, Attila Losonczy

**Author notes:** Correspondence should be addressed to: J.B.P.; A.L.

## Abstract

Neurons in the hippocampus exhibit striking selectivity for specific combinations of sensory features, forming representations which are thought to subserve episodic memory. Even during a completely novel experience, ensembles of hippocampal “place cells” are rapidly configured such that the population sparsely encodes visited locations, stabilizing within minutes of the first exposure to a new environment. What cellular mechanisms enable this fast encoding of experience? Here we leverage virtual reality and large scale neural recordings to dissect the effects of novelty and experience on the dynamics of place field formation. We show that the place fields of many CA1 neurons transiently shift locations and modulate the amplitude of their activity immediately after place field formation, consistent with rapid plasticity mechanisms driven by plateau potentials and somatic burst spiking. These motifs were particularly enriched during initial exploration of a novel context and decayed with experience. Our data suggest that novelty modulates the effective learning rate in CA1, favoring burst-driven field formation to support fast synaptic updating during new experience.

## Introduction

Learning in neuronal systems is complicated by a fundamental tension between stability and plasticity (Carpenter & Grossberg 1991). Networks with fast learning rates encode new information with high fidelity at the expense of overwriting older patterns, while a slow learning rate can preserve existing structure yet frustrate the encoding of novel information. Theoretical studies of neuronal memory capacity suggest that optimal solutions involve concerted processes operating on a spectrum of timescales (Roxin & Fusi 2013, Benna & Fusi 2016). These ideas harmonize with models of multi-stage memory systems in the brain (McClelland et al. 1995): fast-learning circuits can quickly capture detailed memories of new episodes, which are progressively transferred and integrated into slower systems downstream.

The mammalian hippocampus is intimately involved in the formation of episodic memories, and likely mediates an intermediate stage of processing and storage of experiential information prior to long-term storage in the cortex (McClelland et al. 1995). While generally viewed as a short-term memory system, hippocampal dynamics exhibit a diversity of time constants, both at the level of its sub-networks(Mankin et al. 2015, Ziv et al. 2013) and cellular plasticity mechanisms (Bittner et al. 2015, Mehta et al. 1997, Magee & Grienberger 2020). These results are most often derived from the study of “place cells”, excitatory neurons in the hippocampus that are active in specific locations in an environment during exploration (Moser et al. 2008). Spatial behaviors provide a convenient model for studying memory, as the various sensory settings that animals encounter in the environment are organized into highly relational neural representations in the hippocampus (Eichenbaum 2017). Novel population codes develop with remarkable speed, requiring only a few exposures to an environment before a new set of place fields is learned that spans the available space (Wilson & McNaughton 1993, Frank et al. 2004).

A unique synaptic learning rule was recently discovered in the CA1 subregion of the hippocampus, by which pyramidal neurons formed stable place fields within just a few laps after burst firing was recorded from the neuron at a particular location in the environment (Bittner et al.2015, 2017). The sudden emergence of tuning in previously silent neurons marks a rapid reconfiguration of the weights of synapses that were active around the time of the burst event (Bittner et al. 2017, Milstein et al. 2020), with an asymmetric envelope that extends to inputs active several seconds before the event occurred. This “behavioral timescale synaptic plasticity” (BTSP) is a notable departure from conventional plasticity schemes. Its expression depends on the presence of somatic burst firing driven by plateau potentials, which reflect nonlinear input integration in the dendritic arbor of pyramidal cells (Epsztein et al. 2011). These events could be gated by the presence of other factors such as inhibition, neuromodulation or “instructive” inputs signaling reinforcement or novelty (Gerstner et al. 2018, Milstein et al. 2020), which could enable the circuit to rapidly construct new representations during salient experiences.

It remains unknown what connection this plasticity mechanism has to the reorganization of hippocampal responses that occurs when animals are exposed to different environments (“global remapping”, Muller & Kubie (1987)) or in response to salient cues or reinforcement (Hollup et al.2001, Zaremba et al. 2017, Dupret et al. 2010). The presence of plateau potentials or associated somatic burst spiking reliably leads to the formation of place fields *de novo* (Bittner et al. 2015,Diamantaki et al. 2018), but other experiments have reported that new place fields can appear in the absence of these signatures both in familiar and novel environments (Cohen et al. 2017). Clearly the network’s synaptic matrix does not start from a blank slate when learning from each new episode; prior experience in other environments may already provide a weight distribution that produces location-specific, suprathreshold spiking for some neurons, or subthreshold tuning that could be amplified to unmask new receptive fields through other plasticity mechanisms (Lee et al.2012, McKenzie et al. 2021). We lack clarity on the extent to which BTSP contributes to learning hippocampal representations in these different scenarios, which could provide insight on how the network learning rate may change in response to factors such as novelty or salience.

In this work, we conducted a large scale, longitudinal analysis of place field formation during familiar and novel experiences in order to search for signatures of BTSP and how they may change as a function of experience. Using 2-photon functional calcium imaging, we surveyed thousands of place fields and identified an enrichment of BTSP-like dynamics during the initial exposures to a new environment, which then decayed over the course of several days. Our findings suggest that BTSP is a widespread phenomenon in CA1, and illustrate an experience-dependent regulation of plasticity that could be controlled by internal or external factors to dynamically tune the learning rate of hippocampal representations.

## Results

We constructed a virtual reality system for head-restrained mice, comprising 5 LCD displays surrounding a running wheel; movement through the virtual environments was yoked to a rotary encoder on the wheel axle. We combined this apparatus with 2-photon functional calcium imaging in order to record CA1 neural populations as mice explored the virtual contexts (Fig 1A, S1, see Methods for details). Mice were stereotactically injected with an rAAV vector encoding the calcium sensor GCaMP6f under the control of the synapsin promoter, targeted to the CA1 pyramidal layer. We then implanted a chronic window above the hippocampus to provide optical access for imaging experiments (Lovett-Barron et al. 2014). All imaging data was post-processed in Suite2p (Pachitariu et al. 2017) for motion correction, cell detection, and extraction of raw fluorescence traces (Fig 1B). Signals were neuropil-corrected and detrended for baseline drift, and putative events were detected using the OASIS spike inference algorithm (Friedrich et al. 2017). Imaging data was collected from approximately the same area of CA1 for each 3-day experiment sequence, although we did not attempt to track the identity of individual neurons across sessions. The resulting dataset consisted of an average of 391 ±39 CA1 neurons per session (35 sessions from 6 mice).

**Figure 1.**
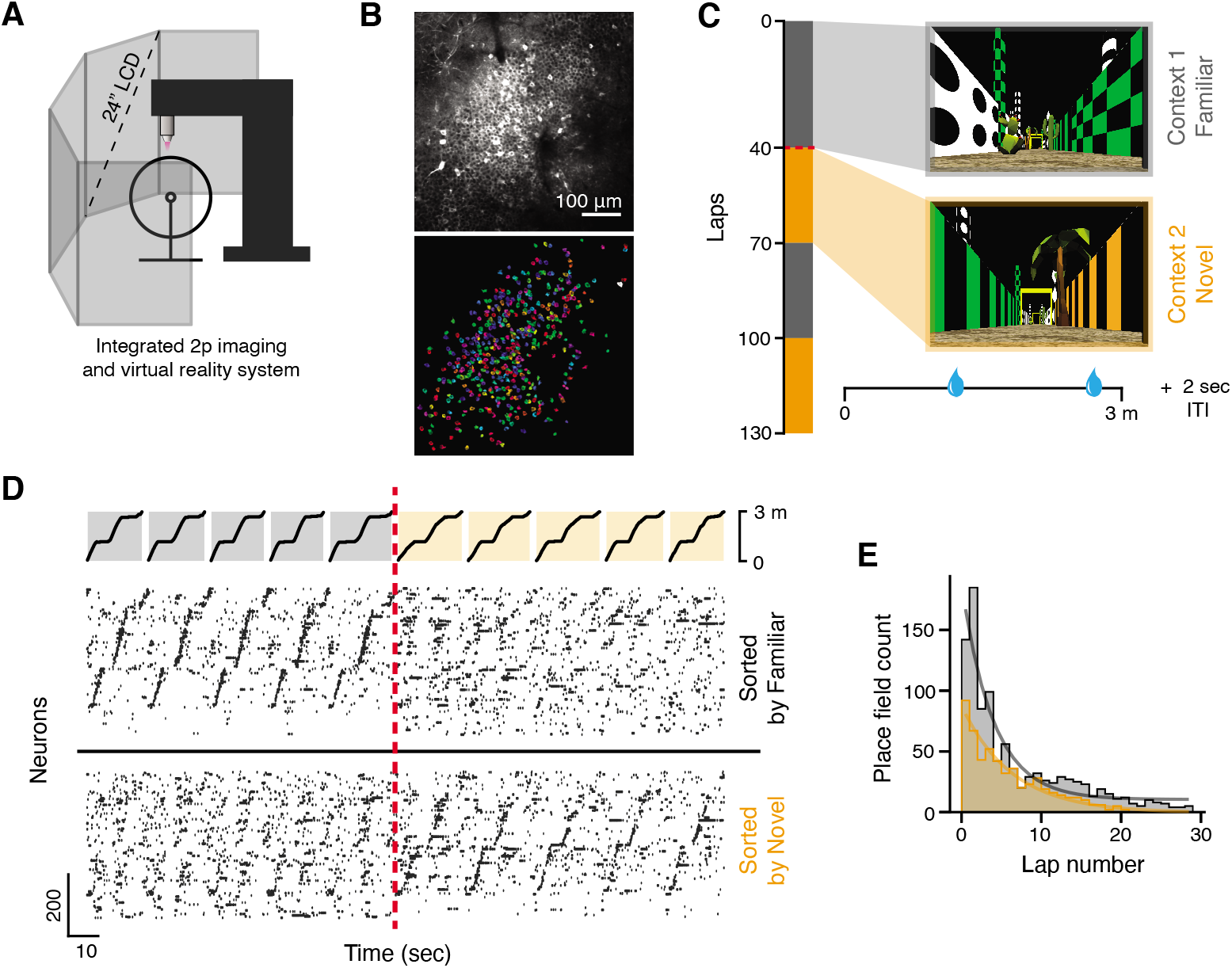
Formation of CA1 representations in novel virtual environments. **A:** Schematic of integrated 2-photon microscope and virtual reality system, for simultaneous *in vivo* functional calcium imaging and head-fixed behavior experiments. **B:** Top: example 2-photon field-of-view (FOV) of CA1 pyramidal neurons expressing the fluorescent calcium reporter GCaMP6f. Bottom: spatial masks of functionally identified ROIs from the same FOV. **C:** Structure of the task. Each session consisted of 130 laps, alternating in blocks between a familiar (grey) or novel (orange) context. Both environments were 3 m in length. Mice were trained to run for sucrose rewards that were delivered at two fixed locations on each lap (~1.2 and 2.7 m); the relative position of reward was the same in both contexts. The experiment was repeated for 3 consecutive days using the same familiar/novel contexts. **D:** Ensemble neural activity from an example Day 1 session, shown for ±5 laps around the very first exposure to the novel context (transition marked in red). Neurons are sorted by their peak firing location on the track, calculated from their average spatial tuning in either familiar (top) or novel (bottom) context laps. A new sequence of place fields rapidly organizes within the first few trials in the novel context. **E:** Place field accumulation in the first trial blocks of the familiar and novel contexts on Day 1, shown as a histogram of the formation laps (the first lap that a cell fired within its place field).

We first trained mice to run for sucrose rewards in a 3 meter virtual environment. All virtual environments began and ended in a darkened tunnel; entering the exit tunnel triggered a 2 sec inter-trial interval (ITI) during which the screens would remain dark, and after which the animal was teleported back to the start of the track. After 1-2 weeks of training, most mice reliably ran over 100 laps in under an hour. To study the dynamics of place field formation during novel experience, we leveraged the virtual reality system to rapidly alternate between familiar and novel environments multiple times during a single experiment (Figure 1C). In each session, mice ran laps through alternating blocks in either a familiar (the training environment) or novel context. Context switches were uncued; the mouse was simply teleported to the other environment at the end of the last ITI of a given block. We repeated the context alternation experiment across 3 consecutive days using the same familiar/novel contexts, to examine the effects of increasing familiarity on CA1 coding. The entire experiment sequence was repeated twice per mouse using two different novel contexts (the familiar context remained the same).

Consistent with previous work using head-fixed VR preparations (Sheffield et al. 2017, Zhao et al. 2020), pyramidal cells’ activity tiled the virtual track, forming a reliable sequence of spatial responses on each lap (Fig 1D). Examining the very first exposure to the novel context on Day 1, a sequence of place fields was rapidly configured in the first few laps following the context switch; the majority of place fields in the novel environment appeared in these early laps, with field accumulation decaying roughly exponentially with experience (Fig 1E) similar to prior reports (Sheffield et al. 2017). Gross place field accumulation in the familiar context was two-fold greater in the initial laps and decayed faster compared to the novel context (*τ* = 3.9 in Familiar and *τ* = 6.4 in Novel for exponential decay fit to Fig 1E). These features agree with our intuition that many place fields in the familiar environment are due to prior learning (and so they immediately appear within the first laps in the session), while the novel representation may continue to grow through ongoing plasticity over a longer time period.

We analyzed the dynamics of place coding across laps in each context block in order to identify activity signatures that may be consistent with different underlying plasticity mechanisms. One hallmark of plateau-induced field formation in CA1 is the backward shift of spatial tuning, relative to the location of burst firing during the formation lap (i.e. the lap when the cell first fires near its place field). This transient shift is a consequence of the asymmetric plasticity kernel of BTSP (Bittner et al. 2017). If new place fields in the novel context form predominantly through BTSP, then this should induce a transient backward drift that is measurable in the population tuning of space during the first few laps in the environment. Since this is the time period during which the majority of place fields form (Fig 1E), the population drift could arise from the cumulative effect of many fields shifting on the same lap, an effect that can notably be measured without first identifying specific place fields or their formation lap.

To quantify any spatial shift between population representations on different trials, we estimated the population spatial cross-correlation between pairs of trials (Figure 2A) by repeatedly computing a population vector correlation as we systematically displaced one trial’s neural responses in space across a range of spatial lags (±75 cm). For the example trial pair in Figure 2A, the peak correlation occurs at a spatial lag of +6.6 cm, indicating that the activity in trial *a* appears forward-shifted in space relative to trial *b*. In Figure 2B, we summarized the resulting spatial shifts between all trial pairs as a matrix, shown for Day 1 experiments (when the animal is exposed to the novel context for the very first time) separately for each context trial block. Since the sign of the the spatial shift between two trials is reversed when the order of comparison is reversed (i.e. trial *a* is after trial *b*, so *b* is before *a*), these matrices are antisymmetric. In general, the upper triangle of the shift matrices was slightly positive, indicating that the population tuning tended to drift backward in space relative to earlier laps in each block (and conversely, the slightly negative lower triangle reflected the tendency of earlier laps to be shifted forward in space relative to later laps). However, the shift pattern during the first exposure to the novel context (Switch 1) was markedly different from other trial blocks: early trials in the context showed a far more exaggerated shift forward in space relative to later trials. We summarized this trend by computing the average pairwise shift for each trial per block (Figures 2C, S2), which showed a large but transient forward shift of population spatial tuning during the early trials of the first block in the novel context. This shift decayed rapidly within the first 10 trials in the new context, notably overlapping with the period during which the majority of new place fields appeared in the novel context (Figure 1E).

**Figure 2.**
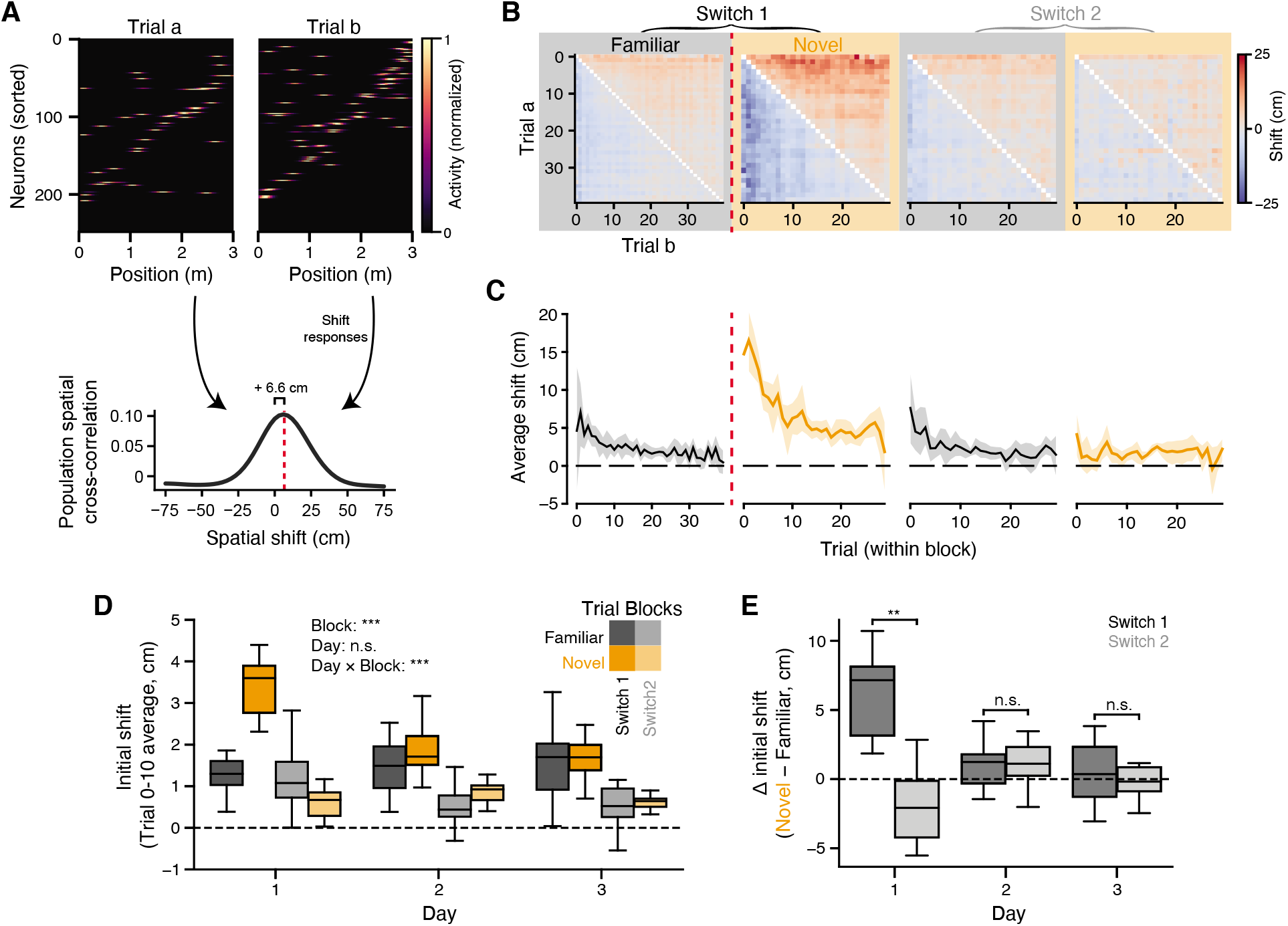
Transient, experience-dependent drift of CA1 ensemble coding. **A:** Top: spatial tuning of the neural population during two example trials in the same context. Neurons are sorted by the location of their peak firing across all laps in the context. Bottom: population spatial-cross correlation computed between the two trials. The correlation at different spatial lags is obtained by shifting each neuron’s tuning in trial *b* before forming the population vector. Red dashed line indicates the center of mass (COM) of the spatial cross-correlation curve. A shift at positive lags indicates the responses in trial *a* are at positions ahead of the responses in trial b. **B:** Spatial shifts (COM) between all pairs of trials during each trial block on Day 0 (average of *n* = 11 experiments). **C:** Average shift for each trial in the experiment, mean and 95% confidence interval across Day 0 experiments. See Figure S2 for individual mice data across all days. **D:** Average representation shifts in the first 10 trials of each block, summarized over all 3 days of the experiment; linear mixed-effects model with main effects of trial block and day (significance shown in inset). **E:** Difference in initial representation drift between the novel vs familiar contexts, for each context switch and day; Wilcoxon signed-rank test. * *p* < 0.05, ** *p* < 0.01, *** *p* < 0.001

The representation in the novel context consistently showed a transient, backward drift during the first exposure across mice on Day 1 (Figures 2D, S3), but this effect was highly experience dependent. We repeated the context switch protocol over a 3 day period, and found that the greatest drift was reliably observed during Day 1, when the animals were exposed to the novel context for the very first time (Figures 2D, E). It is possible that this transient drift is due to the rapid acquisition of new place fields during the initial laps in the novel context. If many neurons acquire their field through BTSP, the population shift can arise as a consequence of averaging over many place fields that shifted acutely after field formation. We simulated this condition and found that it produced a qualitatively similar pattern in the population shift matrices (Figure S3). Other recent reports have suggested that many individual CA1 neurons drift continuously over trials (Dong et al. 2021), but our simulations of this alternative scenario produced population shift patterns that were incongruous with the observed data (Figure S3). Given these comparisons, we reasoned that the population drift was most compatible with frequent BTSP-mediated place field formation in the first exposure to the novel context.

We sought to connect this population-level observation with the behavior of individual place fields (Figure 3A). Due to the asymmetric plasticity kernel of BTSP, individual place fields should acutely shift backward (relative to the direction of animals’ motion) between the formation lap and subsequent laps. We detected place fields in each context block separately for each experiment, and measured how displaced spatial activity on the formation lap was from activity on remaining laps (Figures 3B, C). Note that we use “formation lap” to denote the lap where we first detected stable firing within a place field in a given trial block, but this is not meant to imply that all fields form through ongoing plasticity (i.e. many likely appear simply due to prior learning). Comparing formation lap shifts across different trial blocks on Day 1, we found a clear increase in the displacement of the formation lap’s tuning curve specifically during the first exposure to the novel context, relative to the other trial blocks (Figure 3D). Similar to the population drift, the greatest field shifts were observed during Day 1 on the first context switch (Figure 3E). Examining lap-by-lap displacements of place field activity, we also found that the shifting was most pronounced on the formation lap, and that place fields did not generally continue to drift after the first few trials following place field formation (Figure S4). These results align with the experience-dependent drift in population tuning described in Figure 2, suggesting that the latter arises due to the cumulative effect of many individual cells undergoing acute tuning shifts as they formed their place fields.

**Figure 3.**
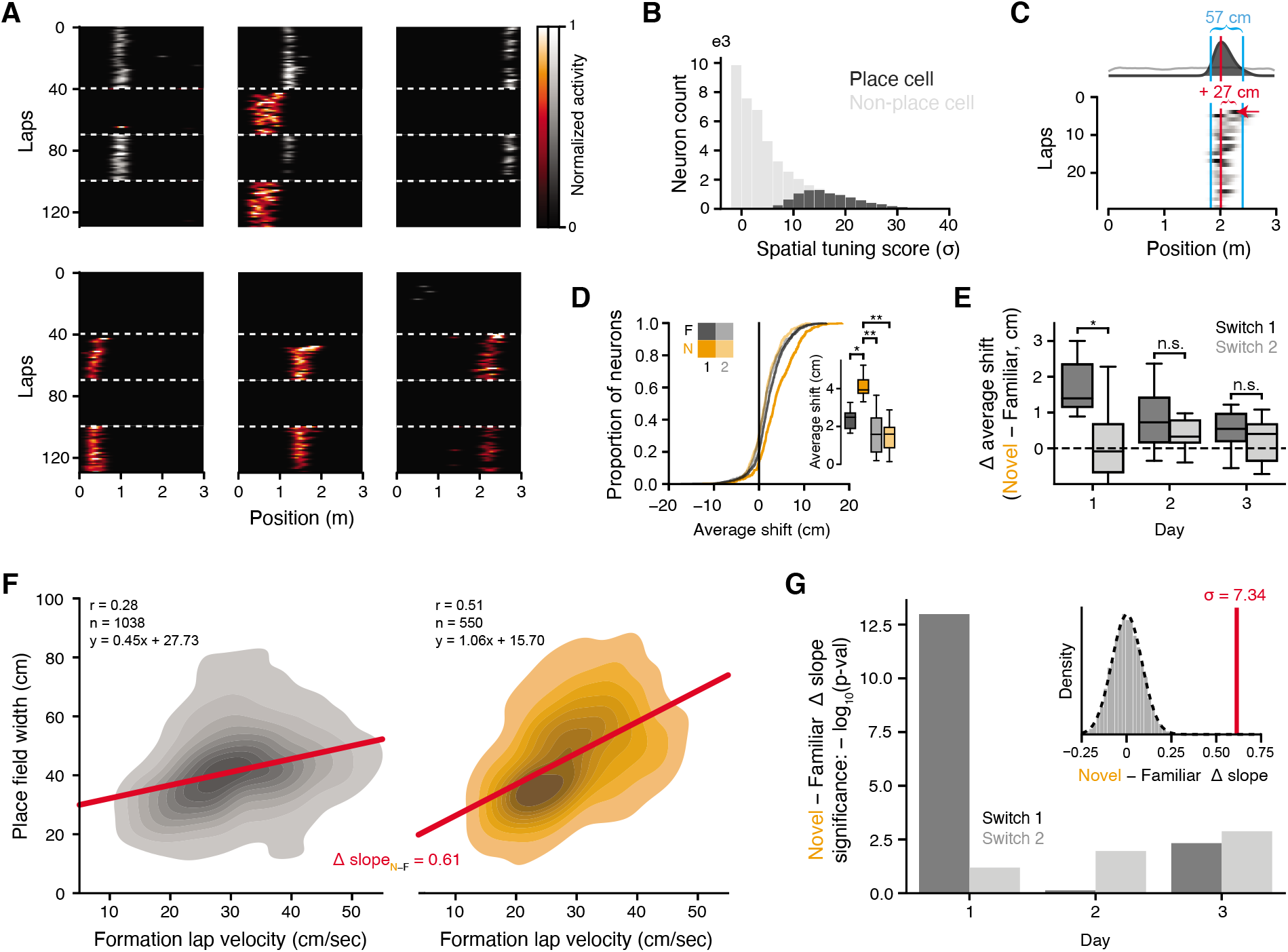
Place field dynamics are consistent with BTSP-mediated field formation in the novel context. **A:** Example simultaneously recorded CA1 neurons with place fields during Day 1 of the experiment. Stable place fields appeared in both the familiar and novel contexts. **B:** Distribution of spatial tuning scores for place and non-place cells. **C:** Characterizations of place field formation for an example neuron: field detection, field width, and formation lap shift. **D:** Distribution of place field shifts (distance between formation lap activity and the remaining in-field activity) during Day 1 of the experiment. Place fields in the first exposure to the novel context exhibit exaggerated spatial shifts on their formation lap. Inset: session-averaged place field shifts on Day 1 (n = 11, Wilcoxon signed-rank test). **E:** Difference in place field shifts between Novel and Familiar contexts, for each context switch and day. The greatest change is seen during the first context switch on Day 1 (n = 11; linear mixed effects model gives significant main effects of Switch (**) and Day (*), and Switch × Day interaction (*); inset: Wilcoxon signed-rank test). **F:** Distribution of place field widths and the velocity of the animal as it traversed the place field during the formation lap on Day 1, switch 1. The linear fit is shown in red. The correlation between velocity and field-width is stronger in the novel context. **G:** Significance of the difference in regression slopes between Familiar and Novel as shown in **F**, for all days and switches. The Δ slope was compared to a null distribution created by randomly permuting the context labels of place fields for each context switch before recomputing the within-context regressions and between-context Δ slope (depicted in the inset for the Day 1, switch 1 results show in **F**). Plotted is the negative log_10_ p-value derived from a Gaussian fitted to the null distribution. Positive values indicate a more significant difference. The strongest difference is seen for Day 1, switch 1. * *p* < 0.05, ** *p* < 0.01, *** *p* < 0.001

The long timescale of BTSP also induces a correlation between the width of place fields and the speed of the animal during the plasticity event: if the animal runs faster, the potentiated inputs will span a larger region on the track (Bittner et al. 2017). For every identified place field, we measured the velocity of the animal as it traversed the place field on the lap of field formation. If many fields during a trial block form through BTSP, then many of these first passes through the place field could contain plateau-driven burst spiking that leads to field formation. Examining the joint distribution of formation lap velocities and the width of the associated place fields, we found that these variables were correlated in both the familiar and novel context (Figure 3F, data shown for Day 1, switch 1). However, this correlation was stronger in the novel context, and the linear fit produced a significantly greater slope compared to the familiar context (Figure 3G). The difference between novel and familiar slopes was also experience-dependent; the first context switch on Day 1 exhibited the most significant slope difference by far across the entire experiment sequence. These results are again compatible with a greater fraction of place cells forming via BTSP during the first exposure to the novel context.

We have so far considered population-level measures that correlate with the relative presence of BTSP-mediated place field formation between experimental conditions. Ideally we would like to segregate individual place fields according to their pattern of spatial drift over laps. By examining groups of place fields that share common spatiotemporal patterns of activity, we could potentially identify a subgroup of place fields exhibiting BTSP-like characteristics or other dynamics, and study additional properties of these classes and how their frequency changes with experience. Toward this aim, we first took an unsupervised approach to identify a set of spatial activity patterns across laps that were shared between place fields (Figures 4A-C). For every place field in the dataset, we examined the first 10 laps of spatial activity, starting from the formation lap. On each of these laps, 150 cm of the spatial tuning curve was extracted, centered around the place field (Figure 4A). We then concatenated all laps from the truncated spatial tuning matrices into a vector for every place field. Consider the example place fields in Figure 4A: since we have centered each field’s activity relative to its field center, the two “flattened” place fields appear as a relatively well-aligned series of bumps. But notably, there is a large deviation in the location of firing during the formation lap: Field 2 exhibits forward shifted activity relative to its place field center, whereas Field 1 remains centered. We hypothesize that many place fields will exhibit similar or other stereotyped patterns of spatial shifts on certain trials. The core idea of the subsequent analysis is to employ dimensionality reduction on these vectorial representations of the place fields, in order to identify a small set of prototypical patterns that summarize the different trends of spatiotemporal activity.

**Figure 4.**
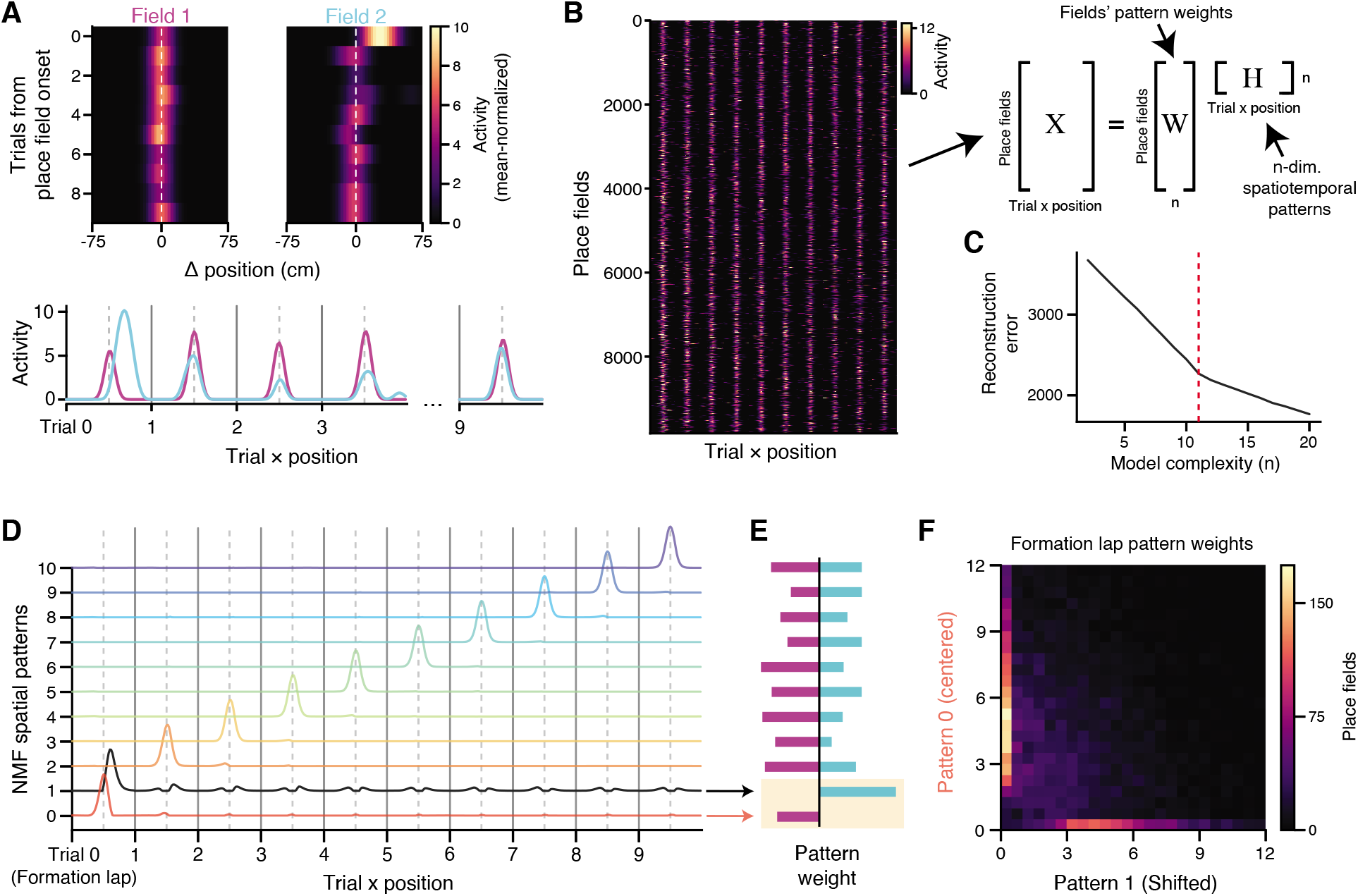
Factorizing place field responses across trials reveals distinct dynamics during the lap of place field appearance. **A:** Top: two example place fields, with data shown in a 150 cm window aligned to the place field center and truncated to the first 10 laps from the onset of place field formation. Bottom: the same fields, “flattened” so that each trial’s spatial tuning curve is sequentially concatenated. **B:** Left: all identified place fields in the dataset with at least 10 laps of activity from onset, cropped and aligned as in **A**. Right: schematic of non-negative matrix factorization (NMF). Each place field’s activity over trials is modeled as a weighted sum (**W**) of a small set of shared spatiotemporal patterns, **H**. **C:** Reconstruction error of **X** for different choices of the number of shared patterns *n* in **H**. There is a clear “elbow” at *n* =11, where model improvement slows. In Fig S6, we also show that this choice of *n* gives the most interpretable patterns. **D:** Spatiotemporal patterns learned by NMF in the *n* = 11 model. The model learns a separate pattern for each lap that produces the centered place field on that lap, plus an additional pattern (in black) on the formation lap that is shifted forward in space relative to the place field center. **E:** Weights of the spatiotemporal patterns in **D** for the example place fields shown in **A**. Trial-to-trial fluctuations in place field amplitude are captured by modulating the weight of the corresponding spatiotemporal pattern. Note the weight differences for the two “formation lap” patterns (yellow shading), which reflect the forwarded shifted activity for the cyan field. **F:** Joint distribution of pattern weights across all place fields for the two “formation lap” components. Field weights for the two patterns are uncorrelated.

All place fields in the dataset were gathered into a matrix **X**, where each row is the spatiotemporal pattern of a single field during the first 10 trials from its formation lap (Figure 4B). Each row was normalized by its mean to encourage the model to focus on shared variability between place fields. We sought a matrix decomposition **X** = **WH**, where each row of **H** would describe a pattern of spatial activity over laps, and each column of **W** would describe how that pattern contributed to individual place fields. There are many techniques for identifying a low-dimensional set of shared patterns in this manner, but here we use non-negative matrix factorization (NMF), since its strict non-negativity aids in interpreting the extracted components (activity described by different components cannot be “cancelled out” due to negative values). In this setting, it is straightforward to interpret the rows of **H** as a pattern of spatial activity over trials, where each individual place field in **X** is modeled as a weighted sum of those different patterns, with the weights given by the rows of **W**. As with any dimensionality reduction, it is necessary to choose the number of components *n* learned by the model. We considered the quality of the reconstruction of **X** across a range of different model complexities (Figure 4C), and identified a clear “elbow” (Milligan & Cooper 1985) between two linear regimes of the reconstruction error at *n* = 11, where adding additional patterns to the model yielded smaller improvements in the fit quality.

Strikingly, the 11-component model results in a very clean and interpretable partitioning of variance in the place field dataset (Figure 4D). 10 of the 11 components encoded the presence of a centered place field on each of the 10 trials included in the data, while the 11th component encoded a forward-shifted place field on the first lap (i.e. the lap of field formation). This is a sensible way to decompose place fields: any individual field can now be reconstructed by appropriately weighting each per-lap component according to the place field’s amplitude on that lap, and the shifted formation lap component can be used to account for variability due to BTSP-like field formation. For example, consider the place fields in Figure 4A. The pattern weights learned from the model accurately reflect the shift in formation lap activity for Field 2, and its reduced amplitude on later laps relative to Field 1 (Figure 4E). Notably, it is not immediately obvious that NMF should find this per-lap representation; the model could instead identify components that are active on several or all laps, representing longer trends of place field amplitude and shifts that are shared between many neurons. Instead, our model’s representation suggests that the variability in place field activity in this dataset is best captured independently lap-by-lap for each place field.

Since the 11-component model converged on this orderly per-lap solution, we wondered if a 10-component model is sufficient to partition the data similarly (since 10 laps are included in **X**), but were surprised to find that it failed to do so. Instead, the 10-component model still prioritized modeling the additional formation lap variance using two components (centered and shifted versions), which resulted in another component encoding a mixture of multiple later trials (Figure S6B). We studied the components learned by models of varying complexity, and found that lower dimensional models invariably resulted in components that encoded mixtures of multiple laps, while adding additional components beyond 11 simply split later lap’s place fields into two smaller fields, symmetrical about the place field center, while still enriching the formation lap representation with additional forward-shifted patterns (Figure S6). Overall, the 11-component model resulted in the most interpretable components, and in light of the clear inflexion point in the loss function (Figure 4C), we focused our remaining analysis on this decomposition.

The representation of formation lap activity in the model illustrates the additional variability present during the lap of place field formation across the dataset. Inspecting the joint distribution of fields’ weights on these two components, we found that they contributed to mostly orthogonal groups of place fields (Figure 4F, note the concentration about the axes). In fact, the two formation lap components had the most orthogonal weight vectors out of any pair in the model (Figure S7A). The shifted formation lap component was also weakly anticorrelated with all remaining lap patterns, which could reflect the tendency for plateau potentials to evoke higher, burst firing rates on the formation lap compared to later laps (this happens because each field is normalized by its mean activity, and so bursting on the first lap will depress the relative amplitude of later lap activity). Given these features, we clustered place fields according to their weights on the 11 NMF components to attempt to isolate a group of fields with BTSP-like characteristics. We found that using two clusters was sufficient to reliably segregate these place fields (Figures S7B-E). In Figure 5A, we plotted the cluster centers for the two groups in the NMF component space. The two clusters are mainly distinguished by their strong, opposing weights on the two formation lap components. Notably, the cluster that highly weighted the shifted “plateau” component also showed lower weights on all subsequent lap components compared to the other cluster. We labeled the clusters as “BTSP-like” (cyan) and “Other” (magenta) based on these features.

**Figure 5.**
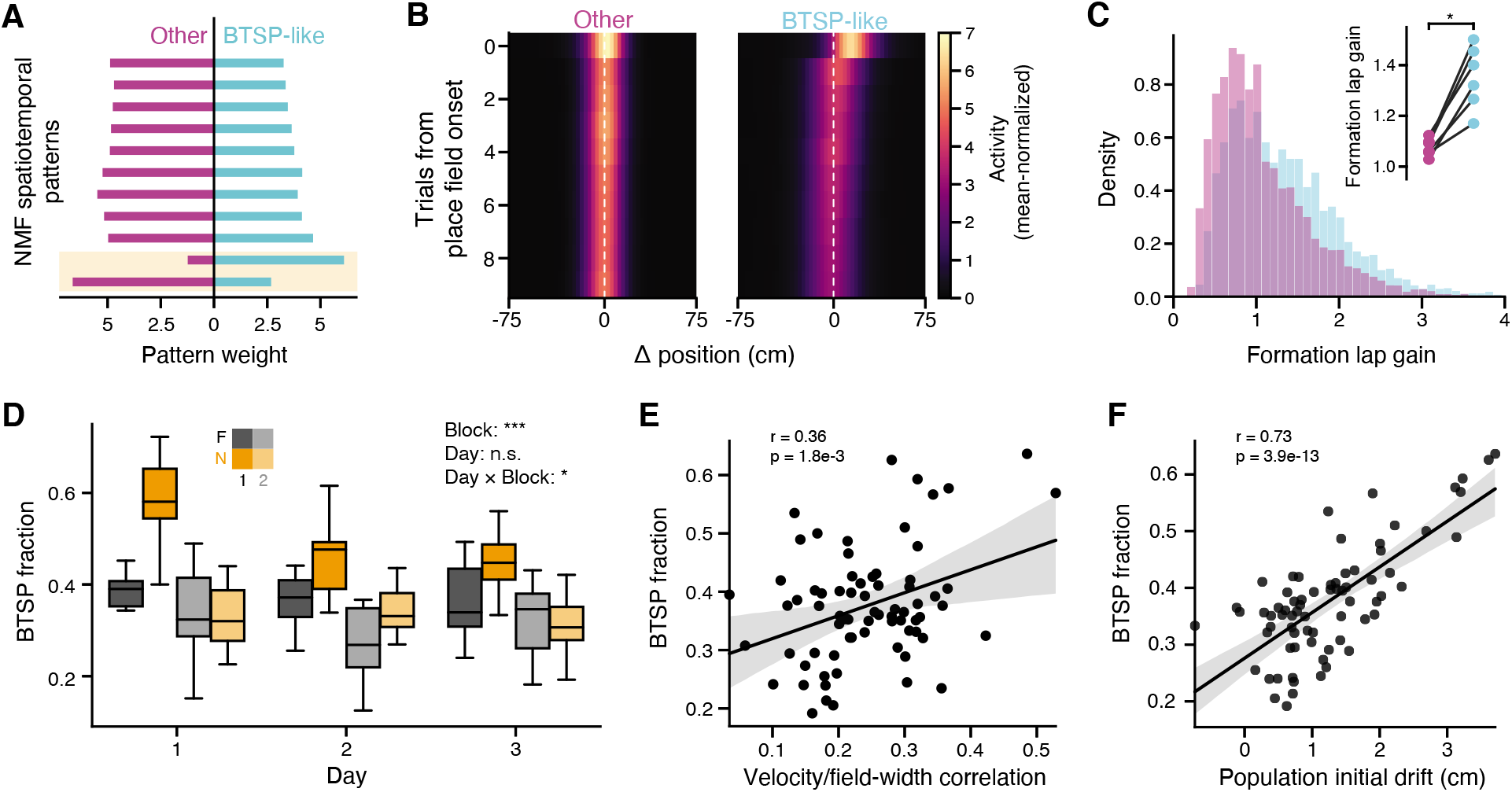
BTSP-like field formation is enriched in the novel context and decays with experience. **A:** Place fields were divided into two groups via K-means clustering in NMF space (i.e., clustering the rows of **W**, see Methods and Figure S7). Plotted is the average NMF pattern weights for place fields in the two groups, labeled as BTSP-like and other. BTSP-like fields have far greater weight on the shifted, plateau-like component on the formation lap, and generally less weight on later lap components compared to the other cluster. Components are sorted as in Figure 4D,E. **B:** Average spatial tuning over trials for place fields in each group. As suggested by the pattern weights in **A**, the BTSP-like group exhibits forward shifted activity on the formation lap, and the formation lap amplitude is greater than later trials. **C:** Formation lap activity gain for each place field, calculated as the peak activity on the formation lap divided by the peak activity from the average of the remaining laps. BTSP-like cells show higher formation lap gain (Kolmogorov-Smirnov test, *p* < 6.68 × 10^−41^). Inset: average gain for place fields in each group, for each mouse (Wilcoxon signed-rank test). **D:** Fraction of place cells classified as BTSP-like in each experiment, shown separately for each trial block on each day. BTSP-like place field formation is enriched during the first exposure to the novel context, and decays with experience. (linear mixed effect model with main effects of trial block and day, significance inset). **E:** Correlation between BTSP fractions and the slope of the velocity/field-width regression. **F:** Correlation between BTSP fraction and the population-level initial drift score. In **E** and **F**, each point is a trial block from a single day and single mouse. * *p* < 0.05, ** *p* < 0.01, *** *p* < 0.001

How do these two groups of place fields differ? We first examined the averaged response profiles for the fields in each group, and found that the BTSP-like fields exhibited strongly forward shifted activity on the formation lap, and relatively reduced amplitude firing on later laps, while the Other fields were well aligned across laps and fired with more consistent amplitude (Figure 5B). We quantified the amplitude change by computing a formation lap gain for each place field, and found that the BTSP-like fields consistently exhibited higher gain (Figure 5C). The two clusters were distinguished by several other features: by construction, the BTSP-like fields exhibited greater shifts in their formation lap activity (Figure S8A), but they also generally exhibited greater place field width on later trials and reduced lap-to-lap stability compared to Other fields (Figures S8B,C). To quantify the isolation of the clusters, we constructed a linear classifier to identify place fields as either BTSP-like or other based on these post hoc characteristics (formation lap gain and shift, field width, and stability) and achieved high accuracy across all mice (85% on average, Figures S8D,E). Further, the clustering identified a highly consistent fraction of place fields as BTSP-like in each mouse (Figure S7C).

Since we could reliably identify a subset of BTSP-like place fields in the dataset, we asked how the fraction of place fields forming with these dynamics changes as a function of experience (Figure 5D). On most days and trial blocks, the fraction of BTSP-like fields was below 40% of all fields, but on the first exposure to the novel context, this increased to nearly 60% on average (Figure 5D). The enrichment of BTSP-like place field formation was also experience dependent, decaying over the course of the three day experiment sequence. The fraction of fields classified as BTSP-like in each condition also correlated with the strength of the correlation between velocity and field width (Figure 5E) and the distance of population tuning drift during the initial trials in each context block (Figure 5F). Overall, our field classification analysis is in good agreement with the population-level measures of BTSP prevalence, and lends further credence to an increased rate of BTSP-mediated acquisition of feature tuning during the initial encounters with novel experiences.

Our data delineated a transient period of enriched, BTSP-like place field formation during novel learning, but it is possible that these dynamics do not affect all components of new experiences homogeneously. Numerous prior works have detailed how hippocampal representations are biased by the presence of salient cues (Bourboulou et al. 2019) and reinforcement (Hollup et al. 2001, Zaremba et al. 2017, Dupret et al. 2010); these effects may be driven in part by the engagement of different plasticity mechanisms. Examining the spatial distribution of place field formation events, we found that BTSP-like and Other fields tended to concentrate in opposing regions of the virtual environments: BTSP-like fields accumulated in the regions between reward zones, while Other fields were particularly enriched near the reward zones (Figure 6A). Notably both of these distributions were correlated with the spatial distribution of velocities (Figure 6B): animals tended to run quickly between reward zones, and reliably decelerated as they approached each reward. It is important to account for this correlation, as our classification of BTSP-like fields is determined principally from the spatial shift in formation lap firing (Figures 4,5,S8E), which will be more difficult to detect when the animal is running at slower velocities.

**Figure 6.**
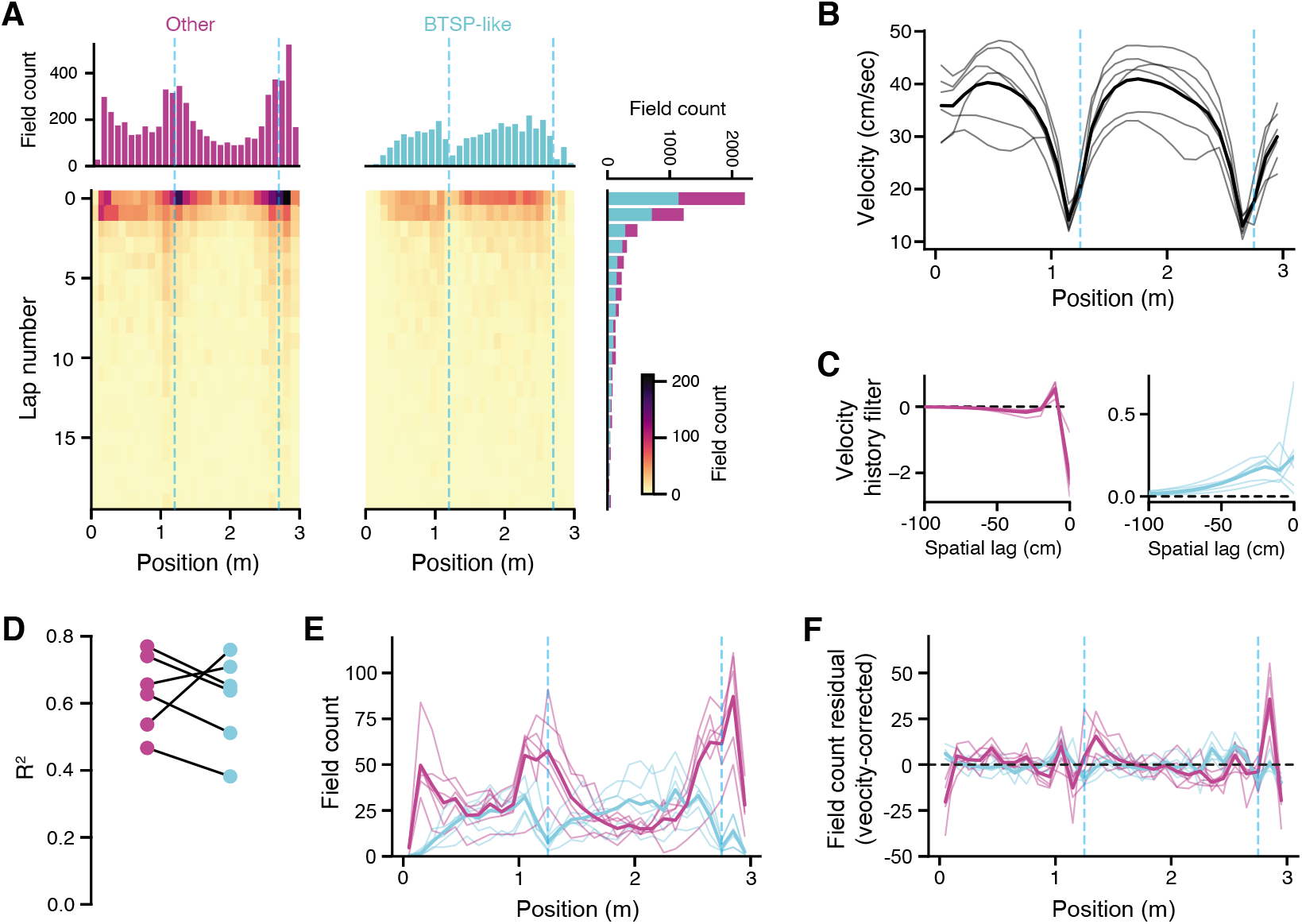
The spatial distribution of place field formation events. **A:** Histogram of place field formation events over positions and lap number (from the start of the context block), with marginal distributions. Data are pooled from all conditions, shown separately for fields classified as BTSP-like and Other in the preceeding analysis. Dashed blue lines indicate reward zones. **B:** Average velocity profile across the environment, for each mouse. The mice slow down considerably on approach to the reward zones. **C:** Spatial kernels fit to predict place field density from the average velocity profile from each mouse. These describe the weighted average of velocity over past and present positions that best predicts the place field density at the present position. Velocity has a large negative effect on Other field density on a short spatial scale. Conversely, BTSP-like field density is positively affected by velocity, on a longer spatial scale. **D:** Prediction quality of the velocity-to-field density model learned in **C**. Filtered velocity explains a large fraction of field density variance for both field types. **E:** As in **A**, the distribution of place field formation events for Other and BTSP-like fields, shown separately for each mouse. **F:** As in **E**, but showing the residual field counts obtained after subtracting off the predictions of the model in **C, D**. The residuals are largely spatially homogeneous and centered at zero, reflecting that the majority of variance is explained by velocity.

We asked how much of the variance in each field formation distribution was explainable by the velocity profile of the animal, by training a linear model to predict field density from the spatially filtered velocity (Figure 6C, see Methods). Our model learned a spatial velocity filter for each animal and each field type. For Other cells, the velocity at the current position consistently predicted a strong negative effect on field density, while for BTSP-like cells, field density was positively predicted by velocity across a range of spatial lags. Overall, the velocity models explained a majority of variance in the distribution of field formation events for both field types (Figure 6D, see also Figure S9). In Figure 6E, we plotted the field distributions for each mouse and field type. Subtracting the model-predicted field distributions from these curves, we found that the residual field counts were largely spatially homogenized and centered at zero, effectively removing the effect of the reward zones (Figure 6F). Given the current data and our field classification methods, we were unable to determine conclusively whether BTSP differentially contributed to the spatial distribution of place fields.

The prior analysis focused on the spatial distribution of place field formation events pooled over all trials. On any given trial though, we hypothesized that the locations of new place fields might be constrained by the representation assembled over prior laps. This could arise from the recruitment of lateral inhibition at locations with existing place fields (Rolotti et al. 2021, Milstein et al. 2020, Robinson et al. 2020), promoting competitive interactions that repel accumulating fields away from one another. The result would be that the spatial distribution of new fields on a given lap is not purely random but is also conditional on the location of recently acquired fields. To test this idea, we examined the formation lap for every place field and measured the average spatial distance between that field and all fields of the same class that formed on the next lap (Figure 7A,B). Since we were interested in the next-lap field distance that was not explained by the overall distribution of place field locations, we normalized this distance by a null distribution that was constructed by randomly permuting the formation lap between fields within a trial block (Figure 7B). In Figure 7C, we plotted the distribution of normalized distances for BTSP and Other fields. For both groups of fields, place fields tended to form farther away from fields that formed on the previous lap than would be expected from random sampling of the environment. This history dependence indicates the presence of competitive interactions that act to disperse new place fields, which could help to drive pattern separation of nearby locations.

**Figure 7.**
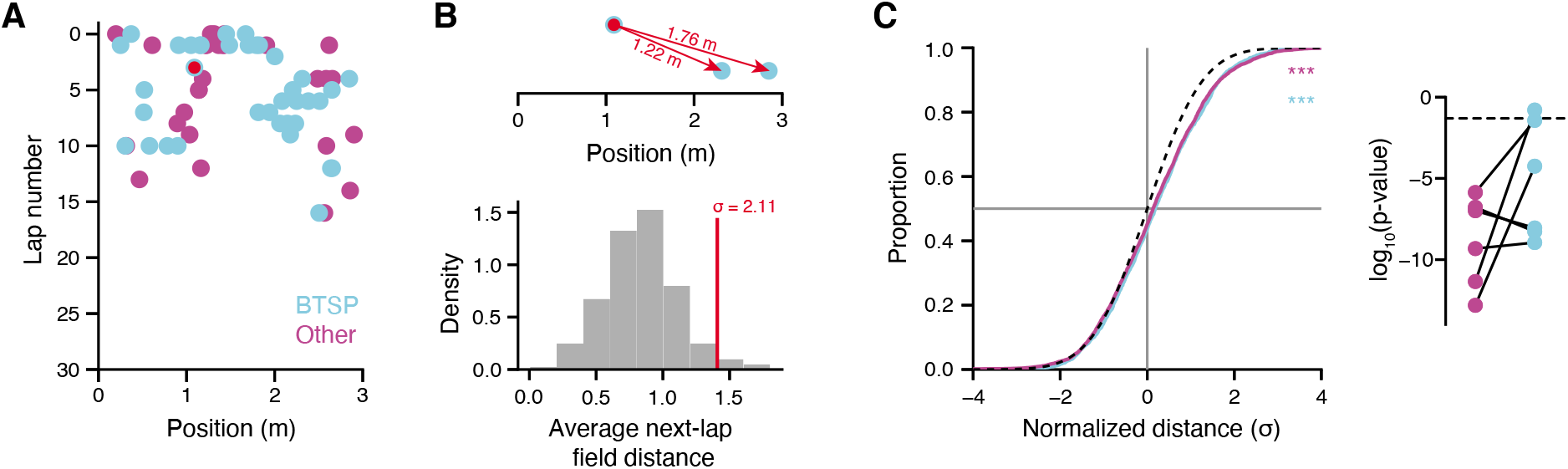
Place field accumulation follows a nonstationary distribution over positions. **A:** Scatter plot of place field formation event over laps and positions, for an example session (Day 1, first novel context exposure). Events are colored according to their BTSP classification. **B:** Illustration of next-lap distance calculation. Top: for every place field, the distance was calculated between its location and the location of all place fields that appeared on the next lap (of the same BTSP classification). Bottom: the average next-lap field distance was compared to a null distribution built by randomly permuting the formation laps of all place fields in the trial block. A normalized distance (*σ*) was derived by standardizing the true distance relative to the null distribution. **C:** Cumulative distribution of normalized distances across the dataset for each place field formation type. The standard normal distribution (chance) is plotted in black. Both BTSP and Other fields show greater next-lap field distances than expected by chance (inset: one-sample Kolmogorov-Smirnov (KS) test against the standard normal distribution). Right: significance of KS test computed for each mouse separately. Place fields accumulated at farther distances compared to prior lap fields than expected by random sampling in nearly all mice and conditions. * *p* < 0.05, ** *p* < 0.01, *** *p* < 0.001

## Discussion

Hippocampal circuits are remarkably plastic, with the ability to construct precise representations of novel experiences with very few exposures (Wilson & McNaughton 1993). This feature is likely critical to the role of the hippocampus in memory storage: it can serve as a fast learning module that rapidly encodes new information, where it is retained and refined for a short period prior to its transfer to and long-term storage in the cortex. (McClelland et al. 1995, Roxin & Fusi 2013). The recent discovery of BTSP is a striking demonstration of the speed of synaptic learning in the hippocampus (Bittner et al. 2015, 2017), and this mechanism could be a crucial component enabling the circuit’s mnemonic functions. In our work here, we have quantified the impact of this novel plasticity rule at the scale of large neuronal populations, and tested a differential role in encoding familiar and novel experiences. Considering multiple lines of analysis, our results are consistent with the hypothesis that a sizable fraction of *de novo* place fields form via BTSP, and that the probability of BTSP events is regulated by the novelty of ongoing experience.

Other reports have questioned the ubiquity of plateau-dependent plasticity events during place field formation (Cohen et al. 2017, Dong et al. 2021). New place fields can appear in a familiar or novel environment in the absence of plateau-associated complex spiking (Cohen et al. 2017), and CA1 neurons can exhibit latent, presynaptic spatial tuning (Lee et al. 2012), which could possibly be amplified to suprathreshold place fields through other synaptic learning processes (McKenzie et al. 2021). In our experiments, we detected signatures of BTSP-like place field formation in all conditions, both at the level of neural ensembles (Figures 2,3) and single neurons (Figures 4,5). While prior work on BTSP used whole cell patch recordings to unambiguously identify plateau potentials, our population-scale approach necessitates that we identify these events through secondary characteristics, namely burst firing and shifted spatial tuning on the first lap of place field appearance. These features are derived respectively from the prolonged somatic depolarization induced by the putative plateau potential (Epsztein et al. 2011, Bittner et al. 2015) and the asymmetric plasticity kernel of BTSP (Bittner et al. 2017), and can be measured through functional calcium imaging at the soma. While these features are indirect, our method allowed us to study place field dynamics at scale and compare the relative frequency of field formation events as animals became increasingly familiar with new environments over days, questions that are impossible to tackle with intracellular recordings limited to single neuron preparations (Bittner et al. 2015, 2017).

All features of BTSP in the dataset exhibited strong experience-dependent effects, with an enrichment of BTSP-related dynamics specifically during the first exposures to the novel environment that decayed over subsequent days. Of course, it is obvious that over multiple days of experience in the same environment, we should expect progressively fewer place fields to appear via ongoing plasticity, with much of the representation simply recalled due to prior learning (although representations continue to drift in familiar environments, see Ziv et al. (2013)). However, our data indicate that burst-dependent plasticity is particularly important during early learning in a novel experience, where our classification estimates that the majority of new place fields appear with BTSP-like characteristics (Figure 5). Plateau potentials are generated through the convergence of coincident presynaptic inputs that triggers active dendritic conductances in the apical tuft (Takahashi & Magee 2009), but the dendritic arbor of CA1 pyramidal neurons is also extensively regulated by local feedback inhibition (Royer et al. 2012, Lovett-Barron et al. 2012). These circuits actively limit dendritic electrogenesis and tightly restrict the number of place fields that can be simultaneously induced via BTSP (Rolotti et al. 2021). This result implies that other factors must be present that transiently free the circuit of these constraints, in order to rapidly build new representations during novel experience.

There is already evidence for a temporary reduction in dendritic inhibition during novel experience in the hippocampus (Sheffield et al. 2017, Geiller et al. 2020). We hypothesize that these inhibitory dynamics are at least partially regulated by factors originating outside of the hippocampus, such as neuromodulatory inputs from the basal forebrain or brainstem (Palacios-Filardo & Mellor 2019) that could be involved in broader, brain-wide signaling of novelty detection. In particular, the locus coeruleus is highly sensitive to novelty and provides dense dopaminergic and noradrenergic input to CA1 (Takeuchi et al. 2016), and these projections have been optogenetically manipulated to increase place field density during a reward learning behavior in mice (Kaufman et al. 2020). Cholinergic neurons are also known to respond to reinforcement and are sensitive to stimulus uncertainty (Hangya et al. 2015), and cholinergic projections from the medial septum ramify extensively in CA1, where they are thought to modulate network function and plasticity in response to arousal (Teles-Grilo Ruivo & Mellor 2013). Characterization of these inputs *in vivo* remains a relatively nascent endeavor, and given the diversity of receptor expression in the hippocampal circuit (Teles-Grilo Ruivo & Mellor 2013), it is likely that they affect both the dendritic excitability of pyramidal neurons and also the recruitment of distinct inhibitory microcircuits. In future work, it will be necessary to assess how neuromodulatory projections affect the generation of plateau potentials and facilitate learning during novel exploration. Overall, we hypothesize that the interplay of external novelty signals and regulation of local inhibition may provide a temporary window of hyper-excitability during new experience, which may permit rapid assembly of representations through BTSP. Abstractly, the temporary shift to frequent burst-mediated place field formation may contribute to an adaptive learning rate in the hippocampus, where the circuit responds to a large change in the input statistics by scaling up the speed of synaptic updates in order to encode new information.

Neuromodulators are often thought to exercise global effects on network state, but other mechanisms may exert more targeted, local influences that act as effectors for BTSP. Specific excitatory inputs may bias the probability of plateau potentials and burst spiking to subgroups of CA1 neurons, either through changes to intrahippocampal afferents (Zhao et al. 2020) or cortical inputs (Boccara et al. 2019, Butler et al. 2019). This upstream reorganization could also occur in response to salience or reinforcement, and serve as an instructive signal for guiding hippocampal plasticity (Milstein et al. 2020). Targeting plasticity to a fraction of the network is one strategy to negotiate the stability-plasticity dilemma and prevent overwriting older memories too quickly.

We found that place fields forming via BTSP were generally less precise than the remaining fields (Figure S8). While many neurons may first reach threshold via BTSP, it seems likely that this initial tuning is refined through secondary mechanisms, such as conventional Hebbian rules and local competitive interactions with other pyramidal neurons mediated through lateral inhibition (Mehta et al. 2000, Cohen et al. 2017, McKenzie et al. 2021, Robinson et al. 2020). Considering that many of the non-BTSP fields are pre-existing place fields learned during previous experience, this could explain why BTSP fields were consistently wider, having not yet undergone further refinement. This is also compatible with the theory that the hippocampus is actively learning a compressed, decorrelated representation of the environment (Gluck & Myers 1993, McClelland et al.1995, Schapiro et al. 2017, Benna & Fusi 2016), as narrower place fields will decrease the correlations between nearby, similar locations. In line with this argument, we also found that the accumulation of place fields was history-dependent (Figure 7). New fields tended to appear at locations farther away from those forming on the previous lap, suggesting that local CA1 networks may facilitate competitive interactions between pyramidal neurons in order to disperse the representation and minimize the neural correlation between nearby locations.

Earlier studies reported that many CA1 place fields undergo a transient period of backward, asymmetric expansion during initial traversals of an environment (Mehta et al. 1997), an effect that encoded the direction of travel and could be modeled by plasticity at feedforward CA3-to-CA1 synapses (Mehta et al. 2000). In our experiments, animals were constrained to run in a single direction through the track, and so it is possible that some component of field width and drift is due to these effects as well, especially residual drift on the trials immediately after place field formation (Figure S4). However, this does not explain the exaggerated first lap shifts and firing rate gains observed in BTSP-like cells nor the enrichment of these patterns during novel exploration, as the asymmetric expansion reported by Mehta et al. (1997) occurred in both familiar and novel environments. Due to the sensitivity of fluorescence calcium indicators, it is also likely that we cannot detect very low firing rate activity and so our ability to resolve any asymmetric expansion is limited. More recently, Dong et al. (2021) reported pervasive and continual backward drift of CA1 place fields in virtual reality. However, the shifts in field locations in our data were overwhelmingly limited to the first few laps following place field formation (Figures 3,4,S4), which agrees with prior work in freely moving animals where individual place fields do not continuously drift (Frank et al. 2004, Mehta et al. 2000).

While here we focused on overall environmental novelty, the hippocampal place code is also heavily influenced by the density and salience of environmental cues (Manns & Eichenbaum 2009,Bourboulou et al. 2019) and the presence of reinforcement (Hollup et al. 2001, Zaremba et al. 2017, Dupret et al. 2010), representations of which may actively shape behavior (Robinson et al. 2020). We searched for evidence of differential concentration of BTSP-like field formation along the virtual track, but found that the distribution was mainly uniform over space after regressing out the component correlated with velocity (Figure 6). In our experiment, the location of reward is highly confounded with the velocity profile of the animal, and so we cannot determine concretely which is the causal factor for the variability in field distributions. Since the field shift associated with BTSP is most apparent at higher running speeds and this is a critical component of our field classification method, we certainly underestimate the rate of BTSP events that occur near the reward zones, and so our results do not preclude a connection between burst-dependent plasticity mechanisms and reinforcement (Milstein et al. 2020).

Our results are congruent with burst-dependent plasticity as an important contributor to representation learning in hippocampal area CA1. It remains unclear whether BTSP is also present in pyramidal neurons in other hippocampal or neocortical networks, and what functional consequences this would have, given the differing circuit architecture (Van Strien et al. 2009). In CA1, pyramidal neurons receive a convergence of inputs from intrahippocampal recurrent networks in CA3, which are believed to store memories through attractor dynamics (Rolls 2007), and from the superficial entorhinal cortex, which can directly relay sensory information about ongoing experience. Novelty detection may be central to the function of CA1 within the broader hippocampal circuit (McClelland et al. 1995, Lisman & Otmakhova 2001). We suggest that the regulation of burst-dependent plasticity in CA1 may selectively permit the integration of novel information into the hippocampus, instigating a cascade of plasticity throughout the hippocampal-cortical loop that could optimize internal representations to adapt to large changes in the statistics of ongoing experience.

## Acknowledgments

We thank Zhenrui Liao, Pamela Rivière, and Nick Robinson for helpful discussions, and for comments on an earlier version of this manuscript. J.B.P. is supported by National Institute of Mental Health (NIMH) F31MH121058. J.C.B. is supported by NIMH F31NS110316. S.V.R. was supported by NIMH F31MH117892. S.F. is supported by DARPA L2M, NSF’s NeuroNex program award DBI-1707398, the Gatsby Charitable Foundation, the Simons Foundation, and the Swartz Foundation. A.L. is supported by NIMH 1R01MH124047 and 1R01MH124867; and National Institute of Neurological Disorders and Stroke (NINDS) 1U19NS104590 and 1U01NS115530.

## Author Contributions

J.B.P., S.F., and A.L. conceived the project. J.B.P. designed experiments, collected data, and performed analysis and modeling. J.C.B. built the virtual reality and behavioral control systems, with assistance from J.B.P. S.V.R. provided critical input on analysis. J.B.P., S.F., and A.L. wrote the paper with input from all authors.

## Methods

### Lead Contact and Materials Availability

Further information and requests for resources and reagents should be directed to the Lead Contact Attila Losonczy. All unique resources generated in this study are available from the Lead Contact with a completed Materials Transfer Agreement.

### Experimental Model and Subject Details

All experiments were conducted in accordance with the NIH guidelines and with the approval of the Columbia University Institutional Animal Care and Use Committee. Experiments were performed with adult (8-16 weeks) male C57Bl/6 mice (Jackson Laboratory).

### Method Details

#### Behavior and Imaging

##### Viruses

Pyramidal cell imaging experiments were performed by injecting a recombinant adeno-associated virus (rAAV) encoding *GCaMP6f* (*rAAV1-Syn-GCaMP6f-WPRE-SV40*, Addgene/Penn Vector Core) into male wild-type mice.

##### Surgical procedure

Methods for viral delivery and surgical implant of imaging window and headposts were largely identical to previous work (Lovett-Barron et al. 2014). Briefly, mice were anesthetized under isofluorane and the virus was injected in dorsal CA1 (−2 mm AP; −1.5 ML, −1.2 DV relative to bregma; 500 nL) using a Nanoject syringe. Mice recovered in their home cage for 3 days following viral infusions. We then aspirated the cortex overlying the left dorsal hippocampus and implanted a 3 mm glass-bottomed stainless steel cannula for imaging access, and cemented a titanium headpost to the skull for head-fixation. For all surgeries, monitoring and analgesia (buprenorphine or meloxicam as needed) was continued for 3 days postoperatively.

##### Behavioral apparatus and virtual reality system

Mice were head-fixed above a low-friction, light weight running wheel (Warren et al. 2021). The axle of the running wheel was coupled to a rotary encoder, which connected to a circuit that decoded and buffered the quadrature data from the encoder and transmitted these position updates to an Intel NUC i5 mini-PC that was used to control the behavior system. As described previously (Kaufman et al. 2020), the experiments were managed through custom software on the PC that would send and receive instructions from a custom GPIO circuit on the behavior apparatus, which managed the water delivery system, lick port sensor and synchronization with the imaging system, as well as from the position tracking circuit and from the virtual reality system. All system elements were connected using high-speed Ethernet and communicated via UDP message passing.

The running wheel was surrounded by 5 LCD computer monitors (Acer SB230 23” IPS screens) arranged in a half-octagon, covering approximately 220°of the visual field of the mouse. Each monitor was connected to an individual ODROID-C2 single board computer running the Android operating system (version 6.0.1) which rendered a fraction of the VR scene on each display. Each ODROID received continuous instructions over Ethernet from the behavior control computer to update the VR environment as the position of the animal advanced. Virtual reality scenes were designed using the Unity game engine.

##### Behavior training

Starting 7 days after implant surgery, mice were habituated to handling and head-fixation as previously described (Lovett-Barron et al. 2014). After two days of acclimation and free running on the wheel, we began exposing the animal to the 3 m virtual environment. The environment used during this training period would later become the “Familiar” context for the main experiment. When the mice reached the end of the virtual track, the screens were momentarily blanked for a 2 sec inter-trial interval, after which they were instantly teleported back to the beginning of the track.

At this point, mice were water deprived to 85-90% of their starting weight. Over a period of 1-2 weeks, we trained mice to run forward through the virtual environment and lick for small volume sucrose solution rewards (5% sucrose, ~ 4*μm* per reward). Rewards were initially dispersed randomly throughout the environment to encourage running, and we slowly reduced the reward count to two fixed reward zones (located at ~ 1.2 and 2.7 m on the track) over the training period. Mice reached training criteria when they could consistently run > 130 laps using the two fixed reward zones in under an hour. Mice were given additional water as needed daily to maintain weights.

##### Familiar-Novel context switching paradigm

For context-switch experiments, mice ran through alternating blocks of trials in the Familiar (training) context and a Novel context that was previously unseen prior to the start of the imaging experiments. Each recording session was organized into 4 blocks: 40 trials in Familiar, 30 trials in Novel, 30 trials in Familiar, and 30 trials in Novel. We additionally repeated this experiment with the same contexts across two additional days, and so a complete experiment sequence represented 3 days of recording. In all trials and contexts, the distance to the sucrose rewards remained fixed at 1.2 and 2.7 m. For each mouse, we also repeated the entire experiment sequence with a second Novel context (the familiar context remained the same for both sequences).

##### 2-photon microscopy

Mice were habituated to the imaging apparatus (e.g. microscope/objective, laser, sounds of resonant scanner and shutters) during the training period. All imaging was conducted using a 2-photon 8 kHz resonant scanner (Bruker) and 16x NIR water immersion objective (Nikon, 0.8 NA, 3 mm working distance). For excitation, we used a 920 nm laser (50-100 mW at objective back aperture, Coherent). Green (GCaMP6f) fluorescence was collected through an emission cube filter set (HQ525/70 m-2p) to a GaAsP photomultiplier tube detector (Hamamatsu, 7422P-40). A custom dual stage preamp (1.4 × 105 dB, Bruker) was used to amplify signals prior to digitization. All experiments were performed at 1.2-2x digital zoom, acquired as 512 × 512 pixels images at 10 Hz.

### Quantification and Statistical Analysis

#### Image preprocessing

Imaging data was organized using the SIMA software package (Kaifosh et al. 2014). Data was motion corrected in Suite2p (Pachitariu et al. 2017) using the non-rigid registration mode. ROIs were also detected using Suite2p (using “sparse mode”), followed by Suite2p’s standard fluorescence extraction and neuropil correction. Identified ROIs were curated post-hoc using Suite2p’s graphical interface to exclude non-somatic components.

#### Neural data analysis

##### Event detection

All fluorescence traces were deconvolved to detect putative spike events, using the OASIS implementation of the fast non-negative deconvolution algorithm (Friedrich et al. 2017). As in Ahmed et al. (2020), we discarded any events whose amplitude was below 4 median absolute deviations of the raw trace, and discretized the resulting signal for all subsequent analysis.

##### Calculating spatial tuning curves

Spatial tuning curves were calculating for each neuron on each lap. The virtual track was discretized into 100 evenly spaced bins (3 cm), which were used to compute a histogram of neural events. After normalizing for animal occupancy, the histogram was convolved with a Gaussian kernel (*σ* = 9 cm) to obtain a smooth activity rate estimate.

##### Population cross-correlation analysis

We used a cross-correlation approach to identify spatial drift in neural tuning across trials, at the level of the neural population. For each pair of trials *a* and *b* in a context block, we can compute their population vector correlation by concatenating the spatial tuning curves of all neurons in the population. Similarly, we obtained the spatial cross-correlation between the two trials across a range of spatial lags by first shifting each neuron’s spatial tuning in trial *b* and then recomputing the population vector correlation with *a*. Here a peak in the spatial cross-correlation at a positive lag indicates that the activity of *a* is ahead of *b* in space (since *b* must be shifted forward to maximize the correlation), and conversely, a peak at a negative lag indicates that activity in *a* is generally at locations behind *b*. We summarized the directionality and distance of the shift between trial pairs by the center-of-mass (COM) of the spatial cross-correlation curve.

The pattern of spatial drift across trials can be visualized by plotting the spatial shifts for all pairs of trials as a matrix. The matrix is necessarily anti-symmetric (since the cross-correlation between *a* and *b* is the mirror image of that between *b* and *a*). Different neural dynamics predict qualitatively different shift matrices: a population of continually drifting place cells would exhibit a diagonally-banded shift matrix (since shift is then a monotonic function of the time between trials), while a more transient population shift (for example, during a period with many place fields forming via BTSP) would appear as vertical/horizontal bands at the affected trials (see Fig. S3 for simulations).

The shifting tendency of each individual trial relative to the whole trial block can be determined by averaging the rows (or columns) of the symmetrized shift matrix (i.e. by multiplying the matrix lower triangle by −1). Symmetrizing is necessary so that all shift comparisons for a given trial consistently indicate the direction of movement. For example, if the neural population is consistently shifting backward, trials before trial *n* will be ahead and trials after will be behind. Averaging the *n*th row of the symmetrized matrix is equivalent to averaging all trial comparisons in the matrix upper triangle that include trial *n*. By our convention, a positive average shift for trial *n* indicates it is part of a generally backward trend, while a negative average shift indicates a forward trend, relative to the surrounding trials in the block. Occasionally activity on some trials correlated poorly with the rest of the block (possibly due to lapses in attention or behavior), and so we excluded any trial pairs where the peak of the spatial cross-correlation was less than 0.1. This method is used to calculate the average shifts in Figure 2C and S?, where we additionally omitted any trials in an experiment where > 1/3rd of trial pair comparisons were missing due to this exclusion criteria.

##### Simulations of place field accumulation

We validated the population cross-correlation on simulated datasets of accumulating place fields, where a sub-fraction of fields exhibited BTSP-like characteristics (first-lap shift and gain), or that exhibited linear drifting over trials. Each simulated population comprised 300 neurons, where each neuron had a 0.2 probability of acquiring a place field. Fields accumulated over 30 laps according to a geometric process with mean 6, which is approximately the observed average first lap for place fields during the first exposure to the novel context during Day 1 of the experiment. Place field centers were sampled uniformly over the environment, and activity was modeled on each lap as a Gaussian bump (*σ* = 5 bins) at the sampled location.

For simulations of BTSP, we fixed a probability of BTSP for place fields in each simulation. If a field was drawn as BTSP, we shifted its first lap activity by a random distance sampled uniformly between 0 and 0.25 of the environment length, and scaled its first lap activity by a random gain sampled uniformly between 1 and 3. We then considered the analysis results for different BTSP probabilities.

For simulation of continuously drifting place fields, we fixed a probability of drifting for place fields in each simulation. If a field was drawn as drifting, we sampled a slope (distance-by-trial) for the drift uniformly from X to X, and shifted each lap’s activity according to that drift function. We then considered the analysis results for different drifting probabilities.

For all neurons, we additionally added out-of-field noise to individual laps with a probability of 0.2. This was simulated as additional Gaussian bumps added to that lap’s activity, centered at a random location in the environment. We additionally applied random scaling noise to each lap, with each lap’s gain drawn 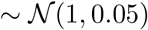.

##### Place field detection

We detected place fields separately in each context trial block (4 per session). Our detection schemed works by finding locations in the virtual environment where a neuron was more active than expected by chance, given its average firing rate and the animal’s spatial occupancy. We constructed a null distribution of spatial tuning curves for every neuron, by circularly shifting each lap’s activity independently by a random distance, and recomputing the smoothed, lap-averaged tuning curve as described previously. This procedure was repeated 1000 times, and we determined the 95th percentile of null tuning values at every spatial bin (i.e. the threshold for *p* < 0.05 spatial tuning). Segments of space where the true spatial tuning curve exceeded this null threshold were marked as candidate place fields, and the place field width was calculated as the distance between where the true tuning curve first exceeded and then again fell below the threshold curve. To restrict our analysis to neurons with unambiguous firing fields, we additionally required that place fields have a width < 1 m (i.e. 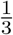 of the track), and that the neuron is active within the bounds of the place field on at least 15 laps.

##### Spatial tuning score

We computed a pseudo-probability mass function *p*(*x_i_*) for each spatial tuning curve, where *x*_1_, *x*_2_, …*x_n_* are the discrete position bins, by normalizing the tuning curve so that it sums to 1 over positions. We then compute the un-normalized tuning score 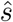 as the KL-divergence between *p* and the uniform distribution *u*:

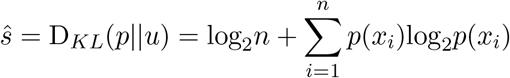

Intuitively, uniform activity over space will give the minimum score *s* = 0, while having all activity concentrated in a single position bin will yield the maximum score of *s* = log_2_(*n*). Since inhomogenous spatial tuning can also arise simply from very sparse, noisy activity, in practice we used a normalized tuning score *s* by standardizing 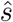 relative to a null distribution, formed by calculating the tuning score on all null tuning curves (generated as described in the place field detection procedure):

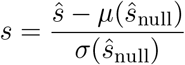

##### Detecting the lap of place field formation

For our analysis of place field characteristics, it was necessary to determine the first lap that a neuron fired within its place field(s) in a given trial block. Following Sheffield et al. (2017), we identified for each place field the first window of 5 laps where the neuron was active within the place field boundaries on at least 3 of those laps, and called the first active lap within that window the place field formation lap. A lap was considered active if there was activity within the place field boundary on that trial with amplitude of at least 5% of the neuron’s peak activity level across all trials. Our results however did not strongly depend on the particular choice of thresholds.

##### Formation lap activity displacement

We estimated the spatial shift between activity on the lap of place field formation and activity on the remaining laps within the place field. This was computed as the difference between the location of peak firing on the first lap, and the location of peak firing in the average tuning of all subsequent laps (as in the population shift analysis, a positive shift indicates the formation lap’s activity was at positions *ahead* of the later lap’s activity).

##### Formation lap gain

Evidence for elevated activity or burst-firing during the formation lap was assessed by computing the formation lap gain, defined as the peak activity rate on the formation lap within the place field divided by the peak activity rate in the averaging tuning of all subsequent laps that contained activity within the place field (i.e. excluding any silent laps, so that the gain does not simply reflect unreliable place fields). A gain > 1 indicates elevated firing during the place field formation lap, relative to later traversals through the place field.

##### Peri-field formation velocity

For each place field, we computed the average velocity of the animal within the place field boundaries during the place field formation lap.

##### Velocity-field width correlation

For each trial block on each day, we fit a linear model to predict the width of place fields from the peri-field formation velocity. We compared the slope of the linear fits between Familiar and Novel contexts by calculating a Δ slope (Novel – Familiar) for each context switch. We calculated the probability that this Δ would be observed under random permutations of the context labels for place fields, recomputing Δ slope for 10000 shuffles in each condition. A p-val was obtained from the cumulative density of the Gaussian fit to the resulting null distribution. We reported the significance relative to the null distribution as the negative log_10_(p-val). In Fig. 3F,G, we pooled place fields across all experiments from a given day and trial block. In Fig. S5, these analyses were repeated for each mouse separately, pooling data from the two repetitions of the experiment sequence for each mouse to reach adequate place field counts.

##### Non-negative matrix factorization

We aimed to identify subsets of place fields in the dataset that exhibited different kinds of trial-to-trial dynamics (e.g. BTSP-like shift in the formation lap activity, drifting responses, or amplitude modulation over trials). To this end, we first aligned all place field tuning profiles by extracting a 150 cm (50 bins) window of activity from each lap around the place field center, and concatenating these windows for the first 10 laps from the formation lap of the place field (e.g. Fig. 4A). This yielded a 500 element vector **x** for every place field, which we stacked into a matrix **X** so that each row was the spatiotemporal profile (trial × position) of a single place field.

Our goal then was to cluster the rows of **X** in order to identify groups of place fields with similar trial-to-trial dynamics, but clustering samples directly in high-dimensional spaces is generally a poor strategy. Instead, we first reduced the dimensionality of **X** using non-negative matrix factorization:

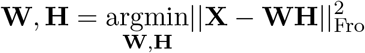

Where **W** and **H** are rank *n* matrices, for some *n* ≪ 500. **H** is an *n* × 500 matrix, where each of the *n* rows is a spatiotemporal pattern. NMF models each row (place field) in **X** as a weighted sum of the these *n* patterns, given by **W**. As with any dimensionality reduction method, since *n* is much less than the number of place fields in the dataset, the model is forced to identify spatiotemporal patterns in **H** that are shared between many place fields. The strict non-negativity of **H** and **W** in NMF however can often give particularly interpretable decompositions, due to its parts-based reconstruction of **X** (i.e., since all elements in the weighted sum for each place field are non-negative, the different patterns in **H** cannot “cancel out”). Each row of **X** was mean-normalized prior to decomposition, so the optimization was not dominated by inhomogenous activity scales across neurons or experiments.

We inspected NMF decompositions of the data for a range of *n*, and found that *n* = 11 corresponded to a prominent “elbow” between two linear regimes in the loss function (i.e. the rate of improvement slows when adding any additional components > 11, Fig. 4C). Additionally, *n* = 11 consistently gave the most interpretable spatiotemporal patterns in **H** (Fig. 4D, S6). In particular, this model produced components that separately represented the place field on each of the 10 laps, plus an additional forward-shifted component on the first lap, reminiscent of plateau-driven place field formation. The weights of the two first-lap components among place fields were strongly anti-correlated (Fig. S6).

##### Clustering place fields in NMF space

We used K-means clustering with *K* = 2 to partition place fields into “BTSP-like” and “other” groups, using the 11 NMF patterns as the feature space for clustering. We found that 2 clusters were sufficient to reliably segregate place fields with BTSP-like characteristics. In particular, we found that increasing the number of clusters had negligible effects on the BTSP-like cluster; additional clusters formed largely through additional subdivisions of the “other” group (Fig S7).

##### BTSP fraction

We computed the fraction of place fields in each experimental condition (day, trial block) that were assigned to the BTSP-like group. In Fig. 4D, we computed this per experimental session. In Fig. 4E,F, this was computed by mouse (pooling each mouse’s data from the two repetitions of the experiment, as was done for Fig S5).

##### Decoding

We validated the NMF-clustering analysis by attempting to decode the BTSP/Other labels of place fields based on secondary characteristics of their activity profiles. We computed 4 features for every place field in the dataset: the shift in its tuning from the first lap to remaining laps, the width of the place field, the stability of its place field (correlation between even and odd laps), and its first lap activity gain. We then trained a support vector machine with a linear kernel to classify place fields as BTSP or Other in this 4-dimensional feature space (Fig. S8). We reported an averaged cross-validated decoding accuracy for each mouse, by randomly partitioning the data into 10 50/50% training/test splits, stratified by BTSP label. Samples were weighted during training to be inversely proportional to label frequency to account for the greater number of Other cells. We additionally compared the cross-validated results to a null distribution constructed by rerunning the cross-validated decoding analysis on copies of the dataset where the BTSP classifications of place fields were randomly permuted. This procedure was repeated 1000 times, and significance thresholds were computed from a 95% interval on the resulting distribution of null accuracies.

##### Modeling the effects of velocity on place field density

We fit a linear model to predict place field density at each position as a function of the animal’s spatial velocity profile. The predictors in the model were exponentially filtered versions of the spatial velocity profile (*τ* ∈ {0.01, 0.025, 0.05, 0.1, 0.25} spatial bins), to account for any short-term history effects, e.g. those induced by the calcium autocorrelation. The velocity filters plotted in Figure 6C are obtained by a weighted sum of the exponential kernels, with the weights given by the regression coefficients.

##### Next-lap field distance

To test for history dependence in the locations of accumulated place fields, we computed for every place field *n* its average distance to place fields that formed on lap *t* + 1, where *t* is the formation lap for field *n*. These distances were computed only between fields of the same BTSP classification. We then standardized this distance according to a null distribution, obtained by randomly permuting the lap of place field formation between all fields of a given BTSP classification within each trial block. In this way, we disrupt the correlations between adjacent laps while maintaining the marginal distributions of place fields over positions and laps. This procedure was repeated 200 times for each place field. The resulting *σ* measures the next-lap field distance relative to the distance expected simply from random sampling according to the marginal distributions.

## Supplementary Materials

**Figure S1.**
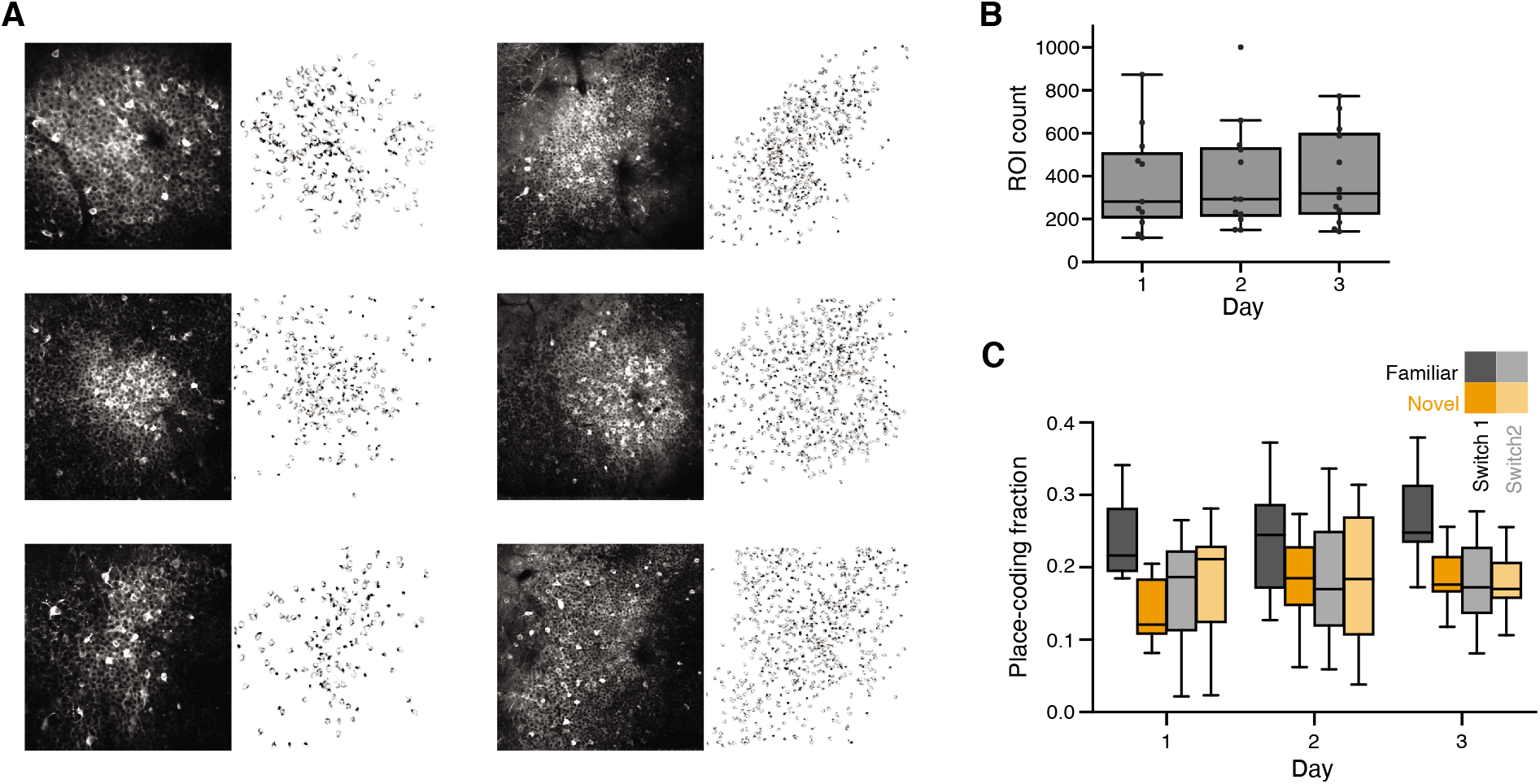
Data survey. **A:** Example FOVs and ROI masks for each mouse in the dataset. **B:** Number of ROIs detected during each experiment, by Day. **C:** Fraction of ROIs with a place field, by Day and trial block.

**Figure S2.**
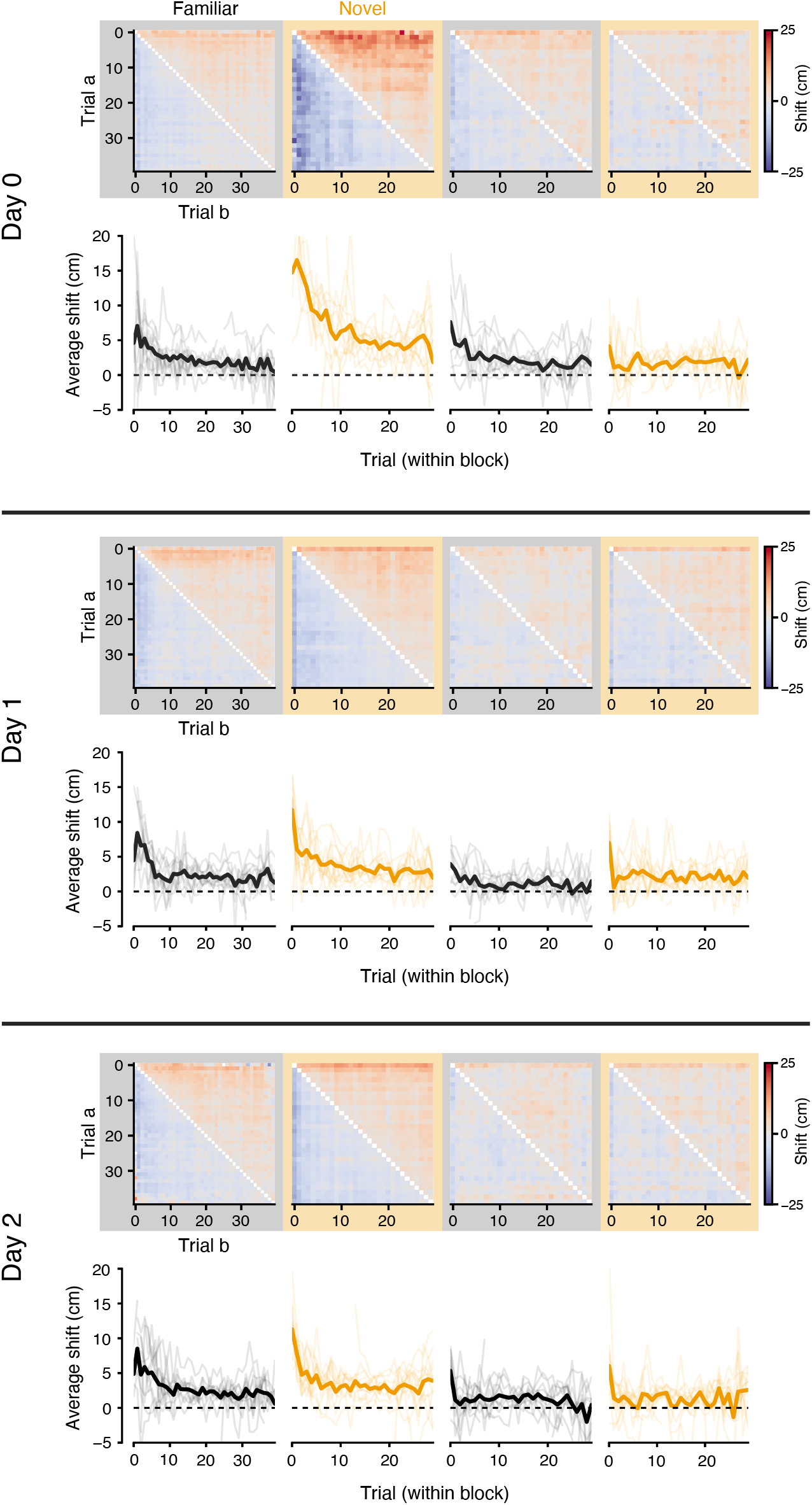
Population spatial drift is transient and experience-dependent. Population drift matrices and average trial shifts, as in Figure 2, shown for all three days of the experiment. The transient, backward shifting representation is specific to the novel context and strongest on the first day of the experiment. These shift distances are summarized and compared across days in Fig 2E, F.

**Figure S3.**
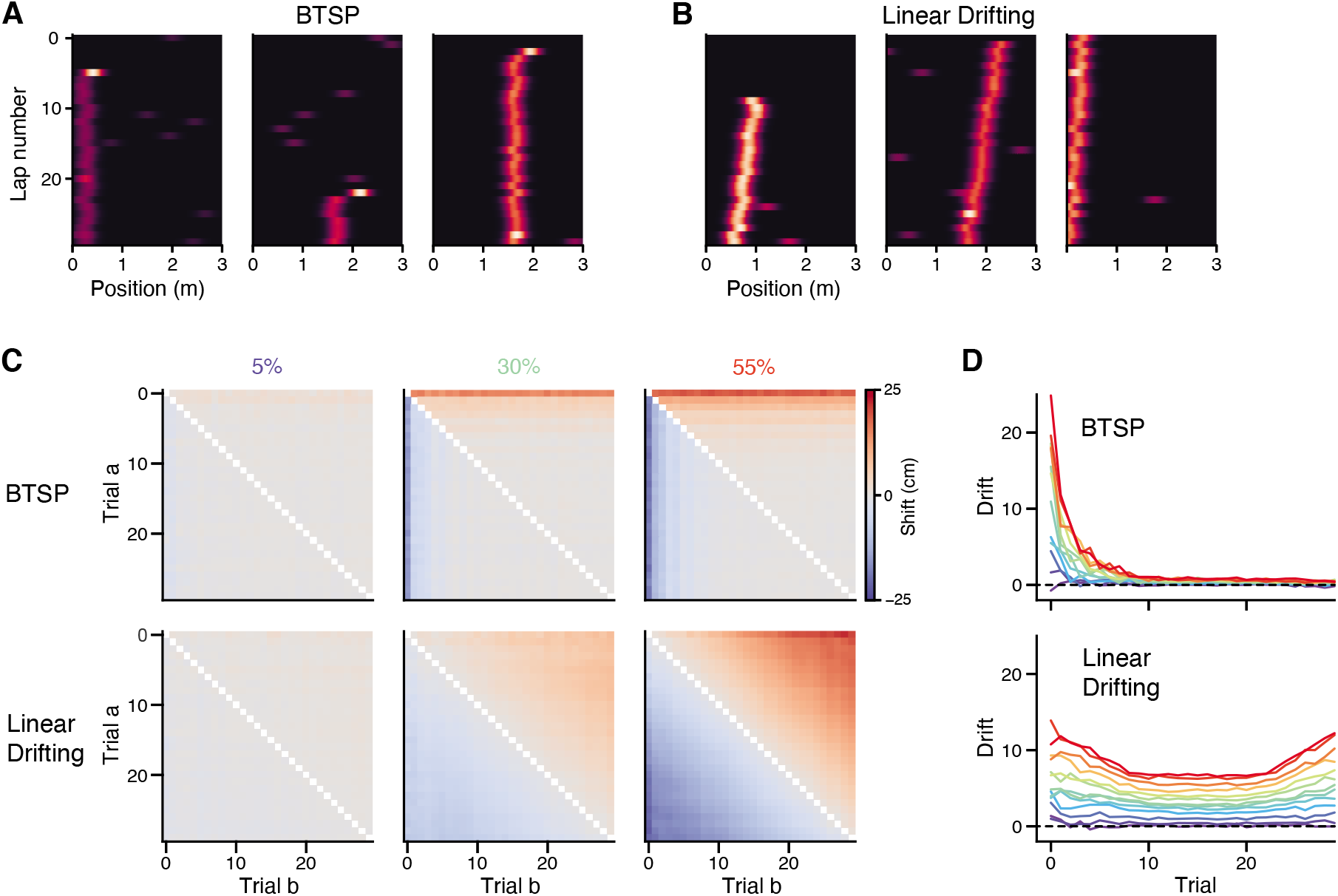
Simulation of BTSP-like vs continuously drifting place fields predict qualitatively different patterns of population tuning drift. **A, B:** Example simulated place fields under the two hypotheses. In both cases, the simulated populations acquired place fields over laps according to a geometric process. For BTSP simulations, each place field had a certain probability of exhibiting BTSP-like characteristics (forward-shifted and gain-modulated first lap activity). For drifting simulations, each place field had a certain probability of exhibiting a linear drift in its field location over laps. **C:** We studied how varying these characteristic probabilities affected the population tuning shift analysis shown in Fig. 2. In BTSP simulations, the population exhibited a transient backward drift concentrated on the first few laps in the session (i.e. when the majority of new place fields form), and this effect became exaggerated with greater BTSP probability. This situation closely resembles the data on the first exposure to the novel context during Day 1, consistent with the population drift arising due to the aggregate effect of many individual neurons undergoing a transient shift on their first active laps due to BTSP. In Linear Drifting simulations, population drift exhibits a very different pattern, where drift is largely a monotonic function of the temporal separation between trials (i.e., each diagonal of the shift matrix is mostly equal). **D:** Shift scores for each trial as in Fig. 2, shown for increasing probabilities of BTSP or linear drifting among the place cell population. Results in **B** and **C** are averages over 20 simulations, where each simulated population contained 300 neurons with a place-coding probability of 20%. See Methods for additional simulation details.

**Figure S4.**
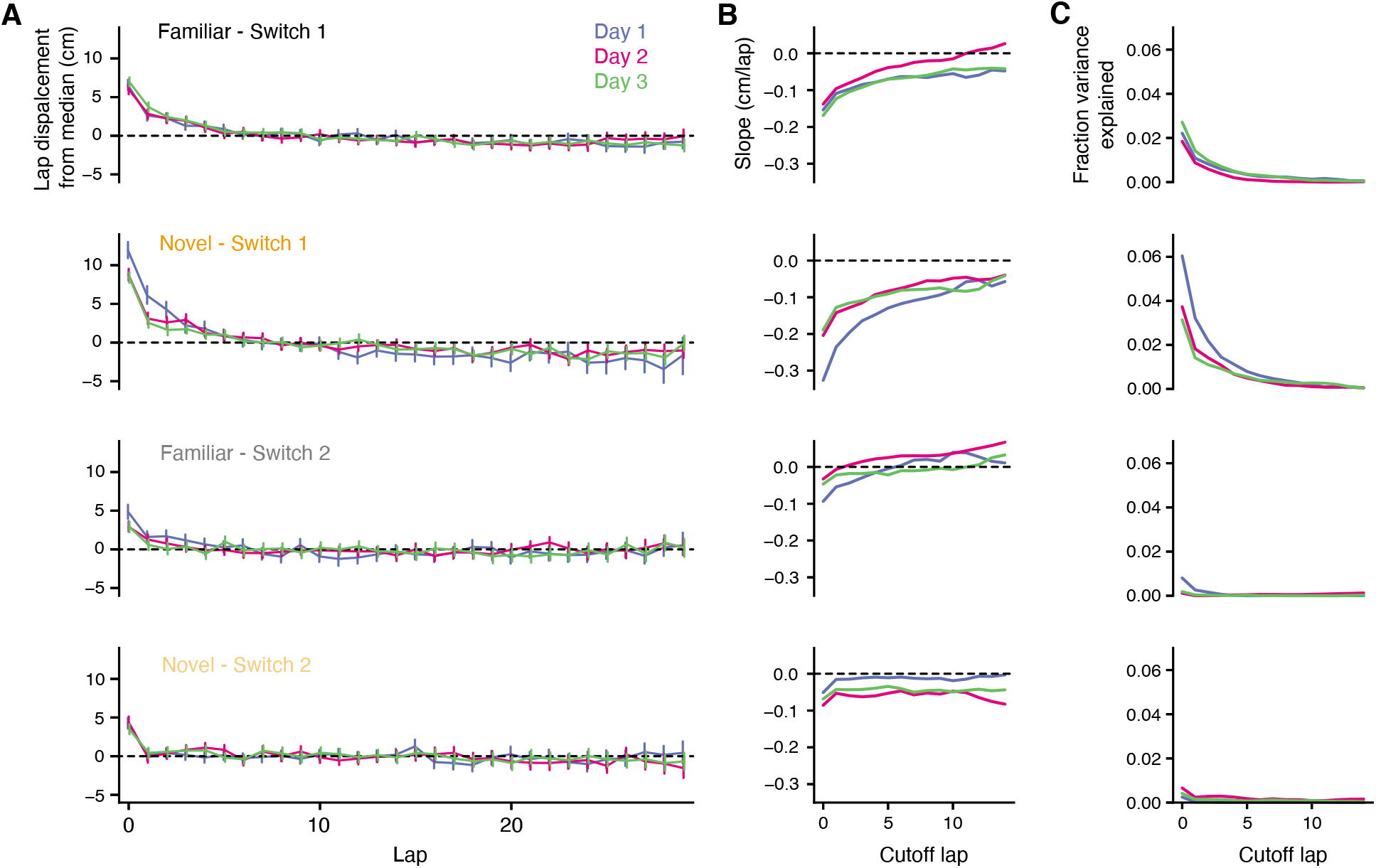
Place fields do not continuously drift in either context. **A:**. Per-lap shifts in the center of mass (COM) of place field tuning, relative to the median lap COM. Place fields are grouped by context trial block and recording day. While there is a transient backward shift present across all conditions in the first few trials of the block, field drift quickly reaches a steady asymptote. **B:** Slope of the linear fit between lap number and field displacement in **A**. We recomputed the regression starting from different laps (‘Cutoff lap’). When initial laps of the trial block are excluded, there linear fit suggests negligible field drift across all conditions (< −0.1 cm/lap, which would accumulate to less than a single discrete spatial bin over the course of a context block). **C:** Fraction of variance explained by the linear fits described in **B**. Excluding the transient drift at the beginning of the trial blocks, the linear trends generally explained < 1% of variance in place field displacements.

**Figure S5.**
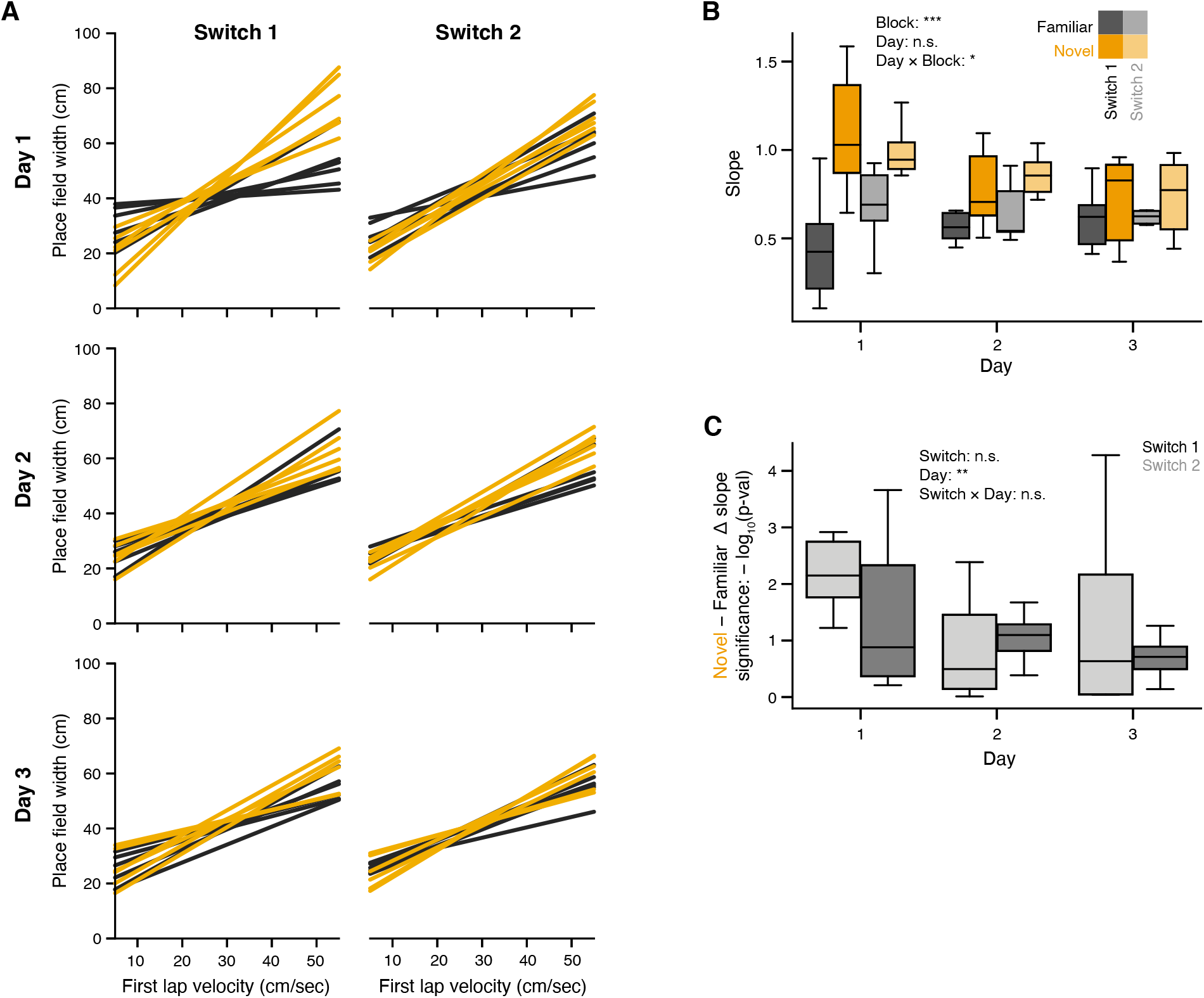
Experience-dependent correlations between field width and first-lap velocity are present in all mice. **A:** Linear fits for place field width as a function of the velocity of the animal during the first traversal of the place field, as in Fig 3F. Regression is computed separate for each mouse, plotted by switch (columns) and days (rows). **B:** Summary of regression slopes in **A**. The slope is greatest for the first exposure to the novel context on Day 1. Linear mixed effects model with main effects of Block and Day, significance inset. **C:** Summary of the significance of the difference between regression slopes in the Familiar vs Novel contexts, as in Fig 3G, computed separately for each mouse, day, and switch. Linear mixed effects model with main effects of Switch and Day, significance inset. * *p* < 0.05, ** *p* < 0.01, *** *p* < 0.001

**Figure S6.**
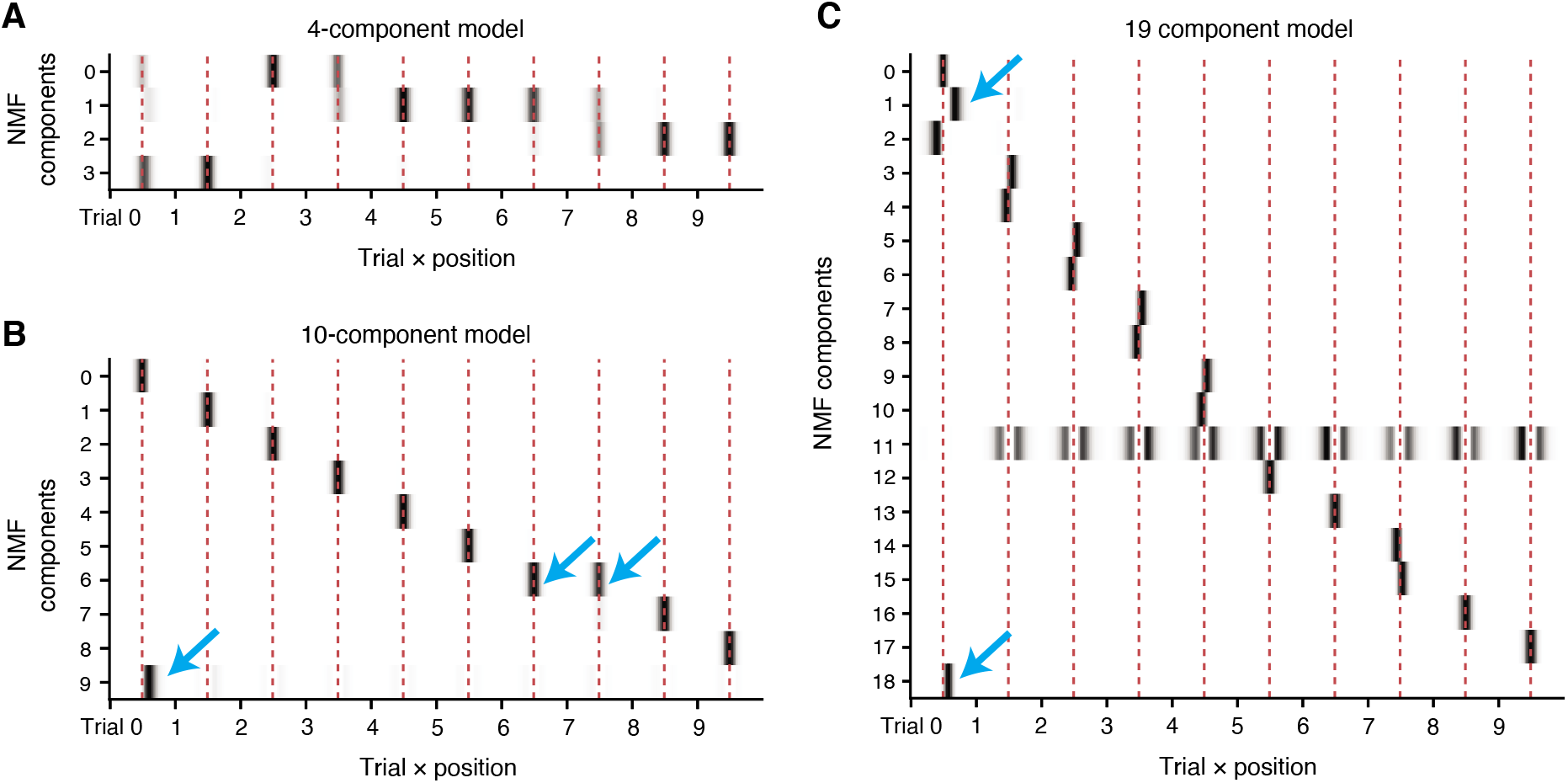
Interpretability of NMF solutions. Example NMF decompositions of the dataset of place fields for different number of components, to illustrate the deleterious behavior of the model for choices other than *n* = 11. Red dashed lines indicate the average lap place field center. **A**: Here *n* = 4 components. We found that when *n* < 10, the components generally ended up reflecting mixtures of adjacent laps. This is likely because, if a place field is active in its field on lap t, it is arguably also likely to be active on laps *t* − 1 and *t* + 1, whereas its activity on more temporally distant laps is perhaps less certain. **B:** Here *n* = 10 components. As *n* increases, more laps obtain a dedicated component for modeling that lap’s place field, but curiously when the number of components is equal to the number of laps, the model still prioritizes modeling the first-lap shift as a separate component. As a result, component 7 subsumes the place field activity of two laps. **C:** Here *n* = 19 components. For all *n* < 11, additional components tended to simply split the later lap place fields in half, symmetrically about the place field center (note how for the laps with two components, the component fields are narrower and lie on either side of the field center). Even still, the model places an outsized importance on capturing the first lap shift in greater detail, indicated by the accumulation of additional components right of the place field center.

**Figure S7.**
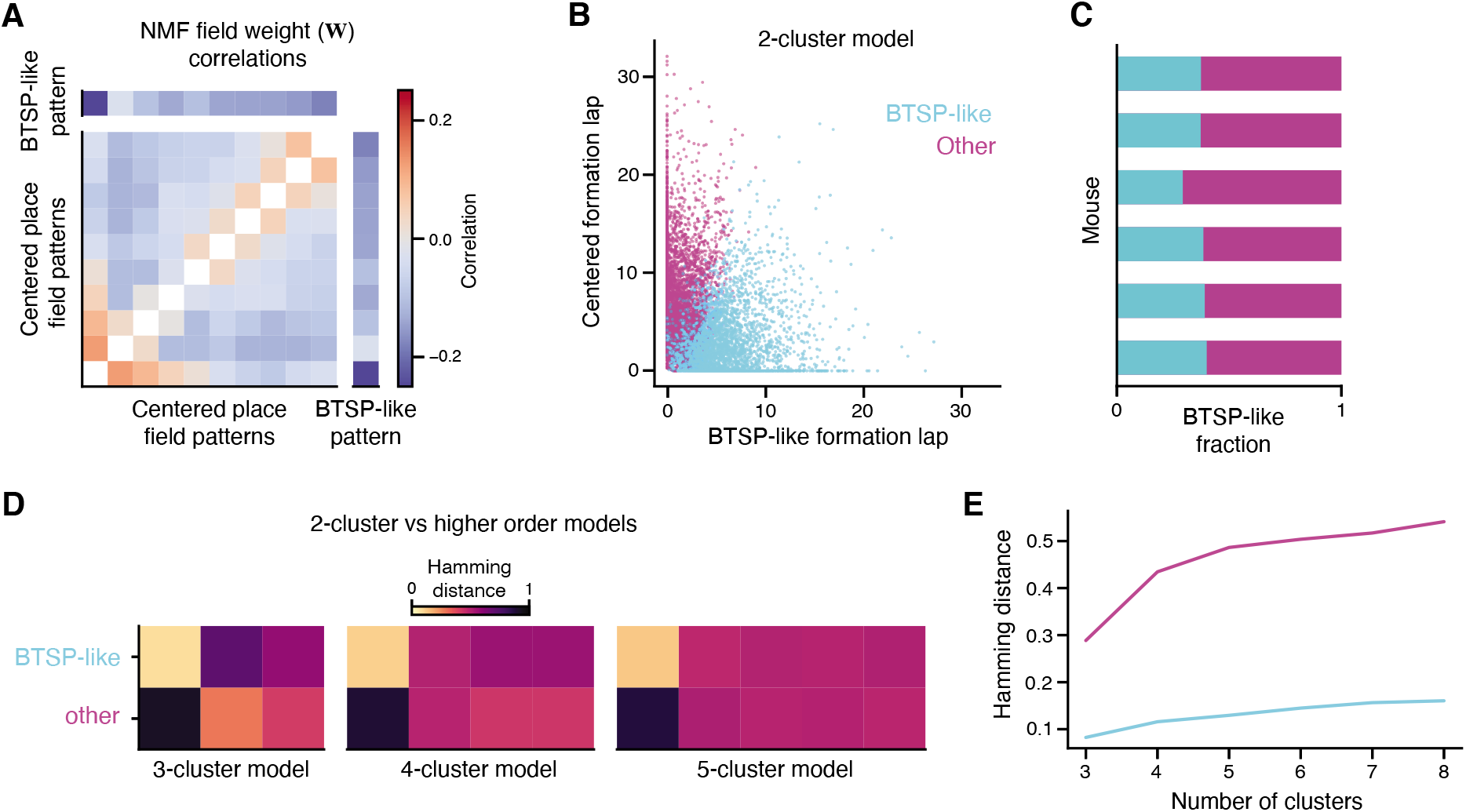
Isolating BTSP-like cells by clustering NMF pattern weights. **A:** Correlation between the place cell weights for each NMF pattern (i.e. correlations between the columns of **W**). The first 10 components are the centered place field patterns for each lap learned by the model, arranged sequentially. The last component is the shifted first lap pattern (”plateau” pattern). The first lap place field pattern and the plateau pattern show the strongest anti-correlation among all pattern pairs, suggesting that differential use of these patterns strongly clusters place fields, in line with the distribution shown in Fig 4F. The plateau component is also weakly anti-correlated with all other lap patterns, consistent with the idea that place fields with a plateau on their first lap will generally have higher amplitude first lap responses compared to later laps. Fields that highly weight the plateau component would then tend to have lower weights for later laps, while fields that down-weight the plateau component will have higher weights for later laps (since each field is mean-normalized across laps). **B:** We applied K-means clustering to place fields in the 11-dimensional NMF space (i.e. clustering the rows of **W**) to identify the BTSP-like group of place fields analyzed in Fig. 5. We set *K* = 2 and found that this was sufficient to reliably uncover a “BTSP-like” group of place fields. Here we show the distribution of place fields in the plane of the centered and shifted first lap patterns, colored by cluster. The BTSP-like cluster subsumes virtually all place fields with weight on the shifted “plateau” pattern. **C:** A similar fraction of place fields are identified as BTSP-like for each mouse in the dataset. **D:** We compared the cluster labels in the 2-cluster model used in the main figures to higher order models. For each order *n* model, we computed the normalized hamming distance between its *n* clusters and the original 2 clusters (BTSP-like/other). A distance of 0 indicates a pair of cluster labelings is identical, whereas 1 indicates they are inverted. Higher order models always find a very similar cluster to the original BTSP-like group, and tend to divide the original “other” group into smaller partitions to populate the additional clusters. **E:** Summary of original BTSP/other cluster survival as the number of clusters is increased. For the BTSP-like and other clusters, we identified the new cluster in each higher order model with the lowest hamming distance (i.e., most similar). The BTSP-like cluster is highly stable.

**Figure S8.**
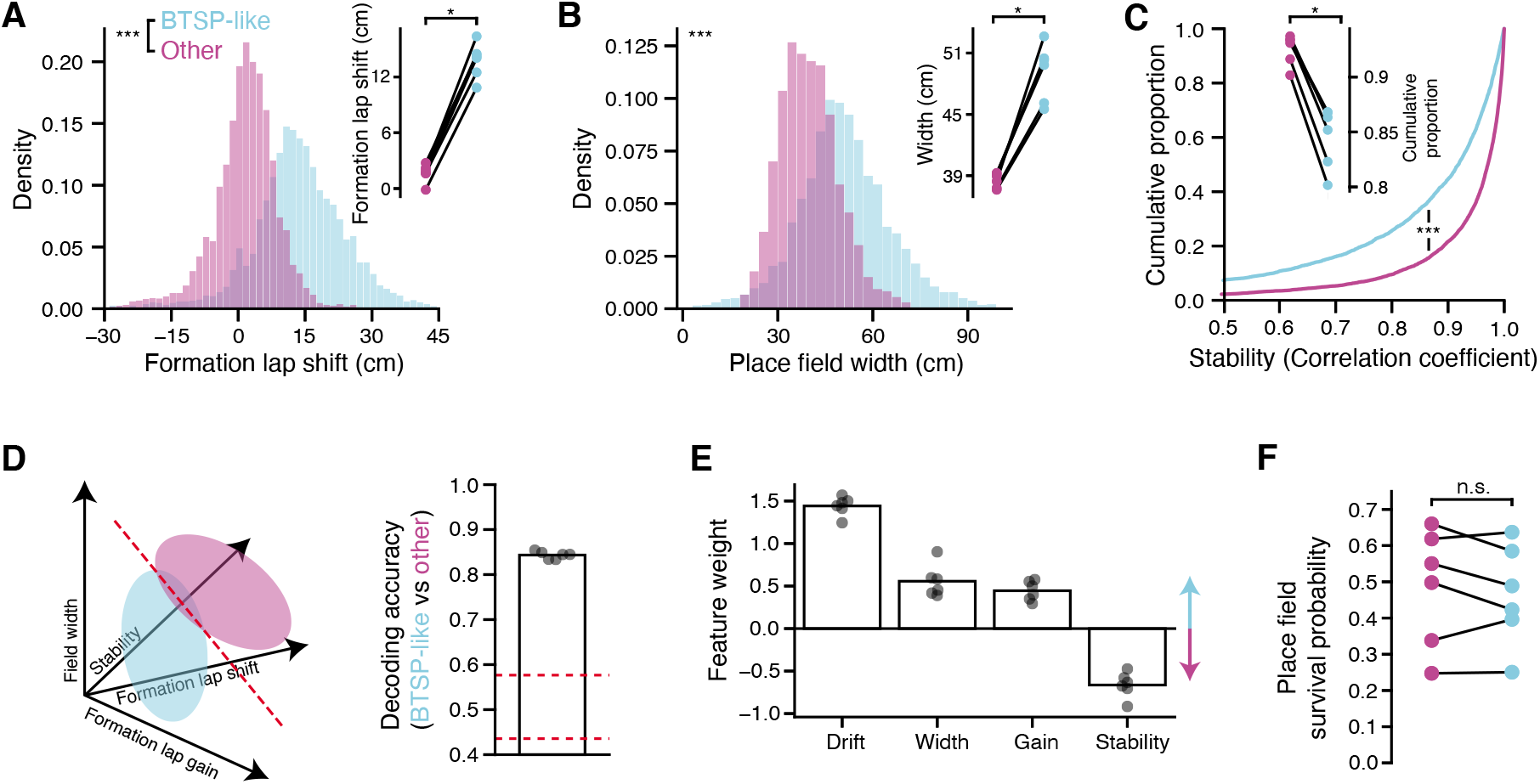
Characteristics of BTSP-like place fields. **A:** Spatial shift between activity on the first lap of place field activity and the activity on remaining laps, for BTSP-like and other place fields. Left: shift distributions for the dataset. Right: average first lap shift for each group, by mouse. **B:** Width of place fields, excluding first lap activity. Plotted as in **A**. **C:** Stability of place fields, computed as the correlation between spatial tuning on even vs odd laps (excluding the first lap). Plotted as in **A**. **D:** Decoding BTSP-like vs Other labels from field characteristics. Left: schematic of the analysis. Each place field was represented as a point in a 4-dimensional space, described by its first lap field gain (Fig. 5), first lap shift, place field width, and stability. A linear SVM was then used to classify fields as either BTSP-like or Other in this space. Right: results of decoding analysis. Plotted is the average cross-validated decoding performance for each mouse. Chance level is demarcated in red, obtained by constructing a null distribution through shuffling place field labels. See Methods for details on cross-validation and significance testing. **E:** Coefficients learned for each feature in the decoder (averaged across cross-validation folds). **F:** Place field survival probability. For each place field identified in the first block of either context, we asked whether we also detected a place field at that location in the second block of the same context during that experiment. Survival probabilities are computed by mouse.

**Figure S9.**
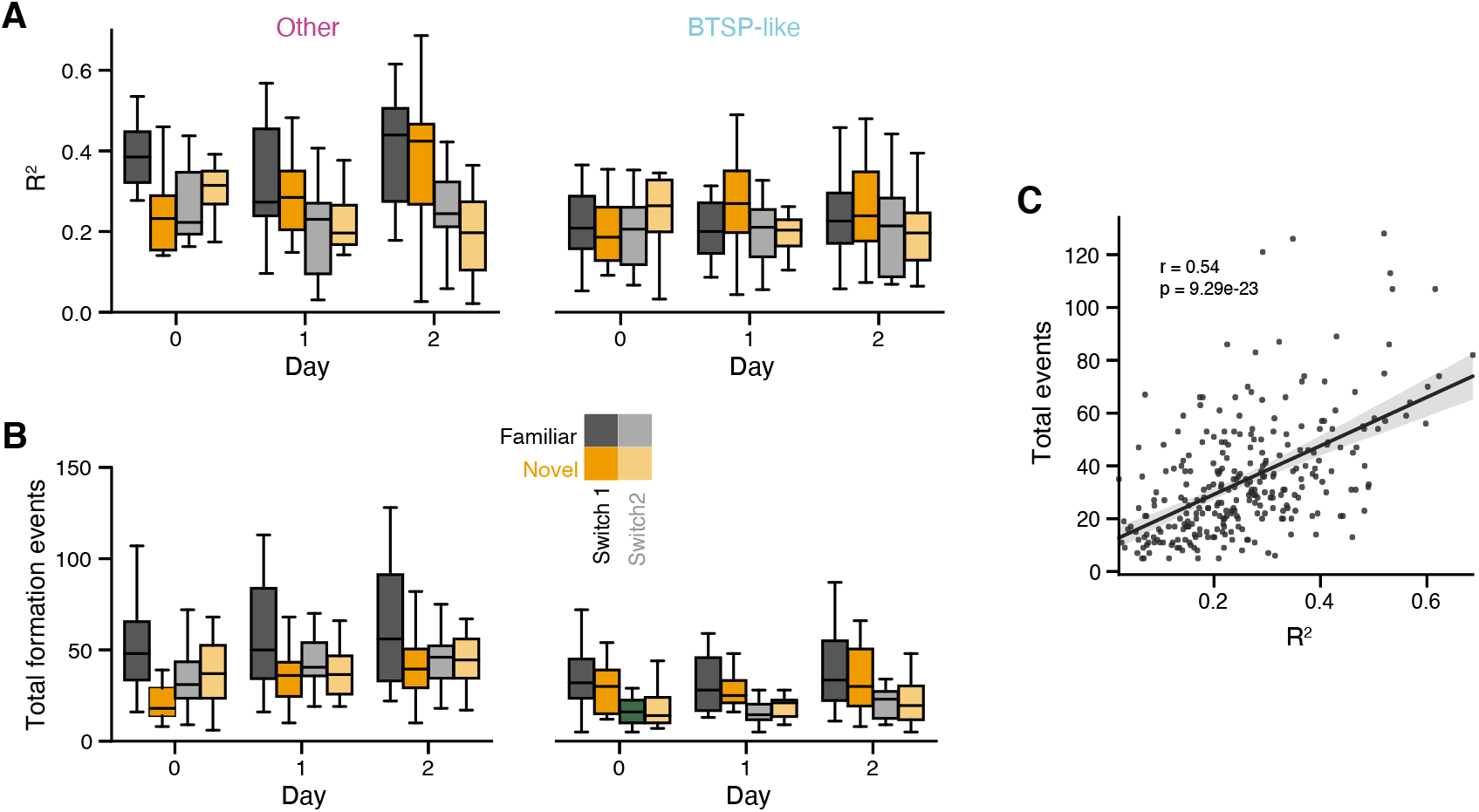
Effect of velocity on the spatial distribution of BTSP across experience. **A:** As in Fig. 6, we fit a linear model that predicted the number of place field formation events at each location as a function of the filtered velocity of the animal. Here we do this analysis separately for each experiment and context block, to identify any changes in the fit quality over experience. Results are plotted separately for ‘BTSP-like’ and ‘Other’ place fields. The quality of the fit varies more for ‘Other’. **B:** Total number of place fields that formed in each of the conditions plotted in **A**. **C:** Correlation between place field counts and the fit quality of the velocity-field density model. Variability in model fit quality is largely accounted for by differences in the number of place field formation events observed during those conditions. Intuitively, if fewer events are sampled in a given condition, it is harder to estimate the true spatial distribution of place field formation.

